# Adaptive laboratory evolution and reverse engineering of single-vitamin prototrophies in *Saccharomyces cerevisiae*

**DOI:** 10.1101/2020.02.12.945287

**Authors:** Thomas Perli, Dewi P.I. Moonen, Marcel van den Broek, Jack T. Pronk, Jean-Marc Daran

## Abstract

Quantitative physiological studies on *Saccharomyces cerevisiae* commonly use synthetic media (SM) that contain a set of water-soluble growth factors that, based on their roles in human nutrition, are referred to as B-vitamins. Previous work demonstrated that, in *S. cerevisiae* CEN.PK113-7D, requirements for biotin could be eliminated by laboratory evolution. In the present study, this laboratory strain was shown to exhibit suboptimal specific growth rates when either inositol, nicotinic acid, pyridoxine, pantothenic acid, *para*-aminobenzoic acid (*p*ABA) or thiamine were omitted from SM. Subsequently, this strain was evolved in parallel serial-transfer experiments for fast aerobic growth on glucose in the absence of individual B-vitamins. In all evolution lines, specific growth rates reached at least 90 % of the growth rate observed in SM supplemented with a complete B-vitamin mixture. Fast growth was already observed after a few transfers on SM without *myo*-inositol, nicotinic acid or *p*ABA. Reaching similar results in SM lacking thiamine, pyridoxine or pantothenate required over 300 generations of selective growth. The genomes of evolved single-colony isolates were re-sequenced and, for each B-vitamin, a subset of non-synonymous mutations associated with fast vitamin-independent growth were selected. These mutations were introduced in a non-evolved reference strain using CRISPR/Cas9-based genome editing. For each B-vitamin, introduction of a small number of mutations sufficed to achieve substantially a increased specific growth rate in non-supplemented SM that represented at least 87% of the specific growth rate observed in fully supplemented complete SM.

**Importance:** Many strains of *Saccharomyces cerevisiae*, a popular platform organism in industrial biotechnology, carry the genetic information required for synthesis of biotin, thiamine, pyridoxine, *para*-aminobenzoic acid, pantothenic acid, nicotinic acid and inositol. However, omission of these B-vitamins typically leads to suboptimal growth. This study demonstrates that, for each individual B-vitamin, it is possible to achieve fast vitamin-independent growth by adaptive laboratory evolution (ALE). Identification of mutations responsible for these fast-growing phenotype by whole-genome sequencing and reverse engineering showed that, for each compound, a small number of mutations sufficed to achieve fast growth in its absence. These results form an important first step towards development of *S. cerevisiae* strains that exhibit fast growth on cheap, fully mineral media that only require complementation with a carbon source, thereby reducing costs, complexity and contamination risks in industrial yeast fermentation processes.

## Introduction

Chemically defined media for cultivation of yeasts (CDMY) are essential for fundamental and applied research. In contrast to complex media, which contain non-defined components such as yeast extract and/or peptone, defined media enable generation of highly reproducible data, independent variation of the concentrations of individual nutrients and, in applied settings, design of balanced media for high-biomass-density cultivation and application of defined nutrient limitation regimes (1, 2). CDMY such as Yeast Nitrogen Base (YNB) and Verduyn medium are widely used in research on *Saccharomyces* yeasts (2, 3). In addition to carbon, nitrogen, phosphorous and sulfur sources and metal salts, these media contain a set of seven growth factors: biotin (B_7_), nicotinic acid (B_3_), inositol (B_8_), pantothenic acid (B_5_), *para*-aminobenzoic acid (*p*ABA) (formerly known as B_10_), pyridoxine (B_6_) and thiamine (B_1_). Based on their water solubility and roles in the human diet, these compounds are all referred to as B-vitamins, but their chemical structures and cellular functions are very different (4). Taking into account their roles in metabolism, they can be divided into three groups i) enzyme co-factors (biotin, pyridoxine, thiamine), ii) precursors for co-factor biosynthesis (nicotinic acid, *p*ABA, pantothenic acid) and iii) inositol, which is a precursor for phosphoinositol and glycosylphosphoinositol anchor proteins (5).

Previous studies demonstrated that growth of *Saccharomyces* species does not strictly depend on addition of all of these B-vitamins, but that omission of individual compounds from CDMY typically results in reduced specific growth rates (6–8). These observations imply that the term ‘vitamin’, which implies a strict nutritional requirement, is in many cases formally incorrect when referring to the role of these compounds in *S. cerevisiae* metabolism (5). In view of its widespread use in yeast physiology, we will nevertheless use it in this paper.

The observation that *Saccharomyces* yeasts can *de novo* synthesize some or all of the ‘B-vitamins’ included in CDMY is consistent with the presence of structural genes encoding the enzymes required for their biosynthesis (Fig. 1, (5)). However, as illustrated by recent studies on biotin requirements of *S. cerevisiae* CEN.PK113-7D (5, 9), a full complement of biosynthetic genes is not necessarily sufficient for fast growth in the absence of an individual vitamin. In the absence of biotin, this grew extremely slowly (µ < 0.01 h^-1^), but fast biotin-independent growth could be obtained through prolonged adaptive laboratory evolution (ALE) in a biotin-free CDMY. Reverse engineering of mutations acquired by evolved strains showed that, along with mutations in the plasma-membrane-transporter genes *TPO1* and *PDR12*, a massive amplification of *BIO1* was crucial for fast biotin-independent growth of evolved strains (10). These results illustrated the power of ALE in optimizing microbial strain performance without *a priori* knowledge of criticial genes or proteins and in unravelling the genetic basis for industrially relevant phenotypes by subsequent whole-genome sequencing and reverse engineering (11, 12).

**Fig. 1:**
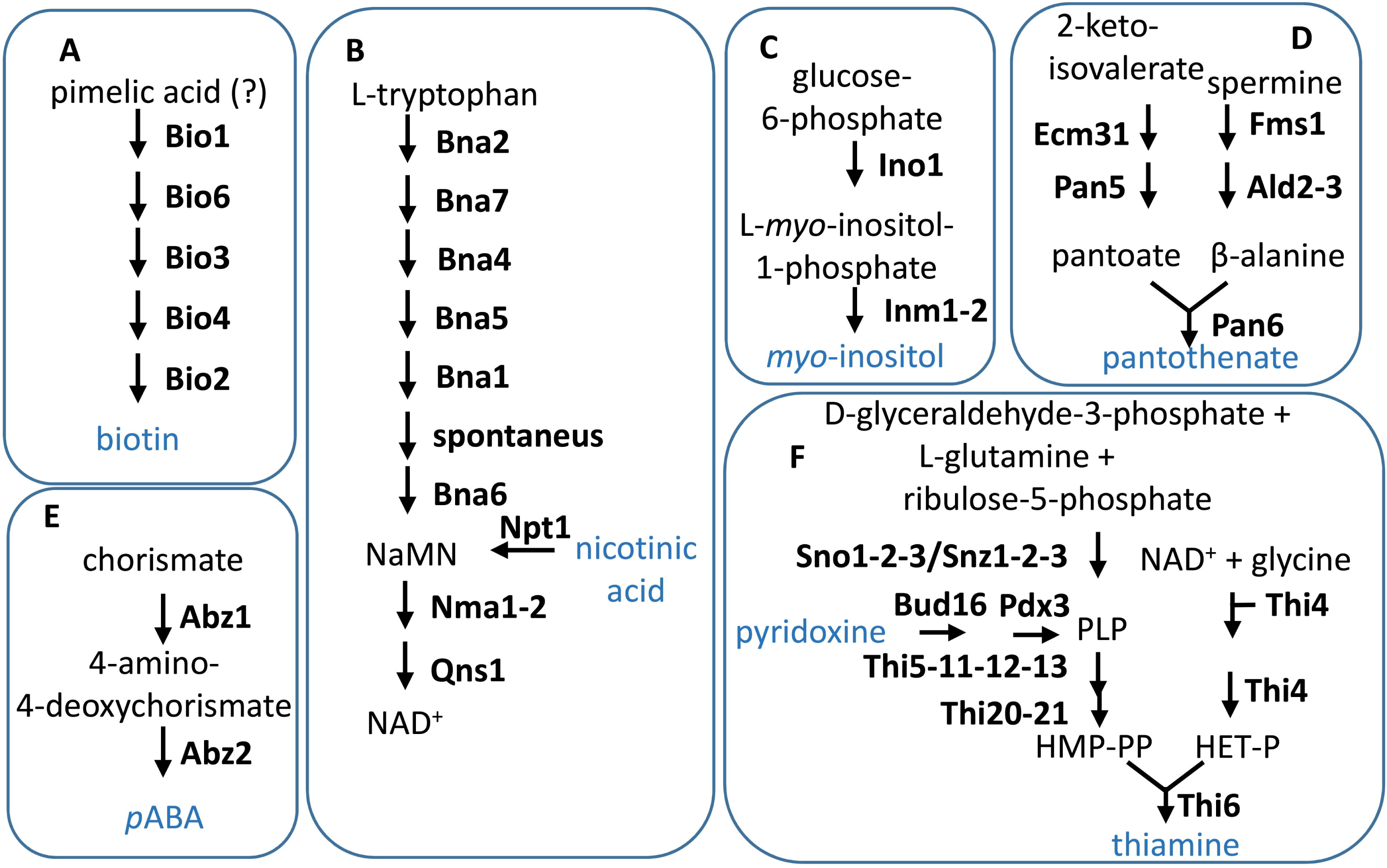
**Schematic representation of the *de novo* biosynthetic pathways for the B-vitamins biotin (A), nicotinic acid (B), myo-inositol (C), pantothenate (D), pABA (E), pyridoxine and thiamine (F) in *S. cerevisiae*** (5). Vitamins that are usually added to the chemical defined media for cultivation of yeasts are shown in blue.

Elimination of vitamin requirements could enable cost reduction in the preparation of defined industrial media and fully prototrophic strains could provide advantages in processes based on feedstocks whose preparation requires heating and/or acid-treatment steps (e.g lignocellulosic hydrolysates; (13, 14)) that inactivate specific vitamins. In addition, processes based on vitamin-independent yeast strains may be less susceptible to contamination by vitamin-auxotrophic microorganisms such as lactic acid bacteria) (15). Thus, chassis strains able to grow fast in the absence of single or multiple vitamins would be therefore be of interest for industrial application. Moreover, engineering strategies aimed at enabling fast growth and product formation in the absence of single or multiple vitamins may be relevant for large-scale industrial application of *Saccharomyces* yeasts.

The goals of the present study were to investigate whether full single-vitamin prototrophy of *S. cerevisiae* for inositol, nicotinic acid, pantothenic acid, *p*ABA, pyridoxine or thiamine can be achieved by ALE and to identify mutations that support fast growth in the absence of each of these vitamins. To this end, the laboratory strain *S. cerevisiae* CEN.PK113-7D was subjected to parallel aerobic ALE experiments that encompassed serial transfer in different synthetic media, which each lacked a single B-vitamin. Independently evolved strains from each medium type were then characterized by whole-genome resequencing and the relevance of selected identified mutations was assessed by their reverse engineering in the parental non-evolved strain.

## Results

### Assessment of CEN.PK113-7D specific B-vitamin requirements

*S. cerevisiae* strains belonging to the CEN.PK lineage, which was developed in an interdisciplinary project supported by the German Volkswagen Stiftung between 1993 and 1994 (16), exhibit properties that make them good laboratory models for yeast biotechnology (17). To provide a baseline for ALE experiments, specific growth rates of the haploid strain CEN.PK113-7D were analysed in aerobic batch cultures on complete SMD and on seven ‘SMDΔ’ media from which either biotin, inositol, nicotinic acid, pantothenic acid, *p*ABA, pyridoxine or thiamine was omitted. To limit interference by carry-over of vitamins from precultures, specific growth rates were measured after a third consecutive transfer on each medium (Fig. 2A).

**Fig. 2:**
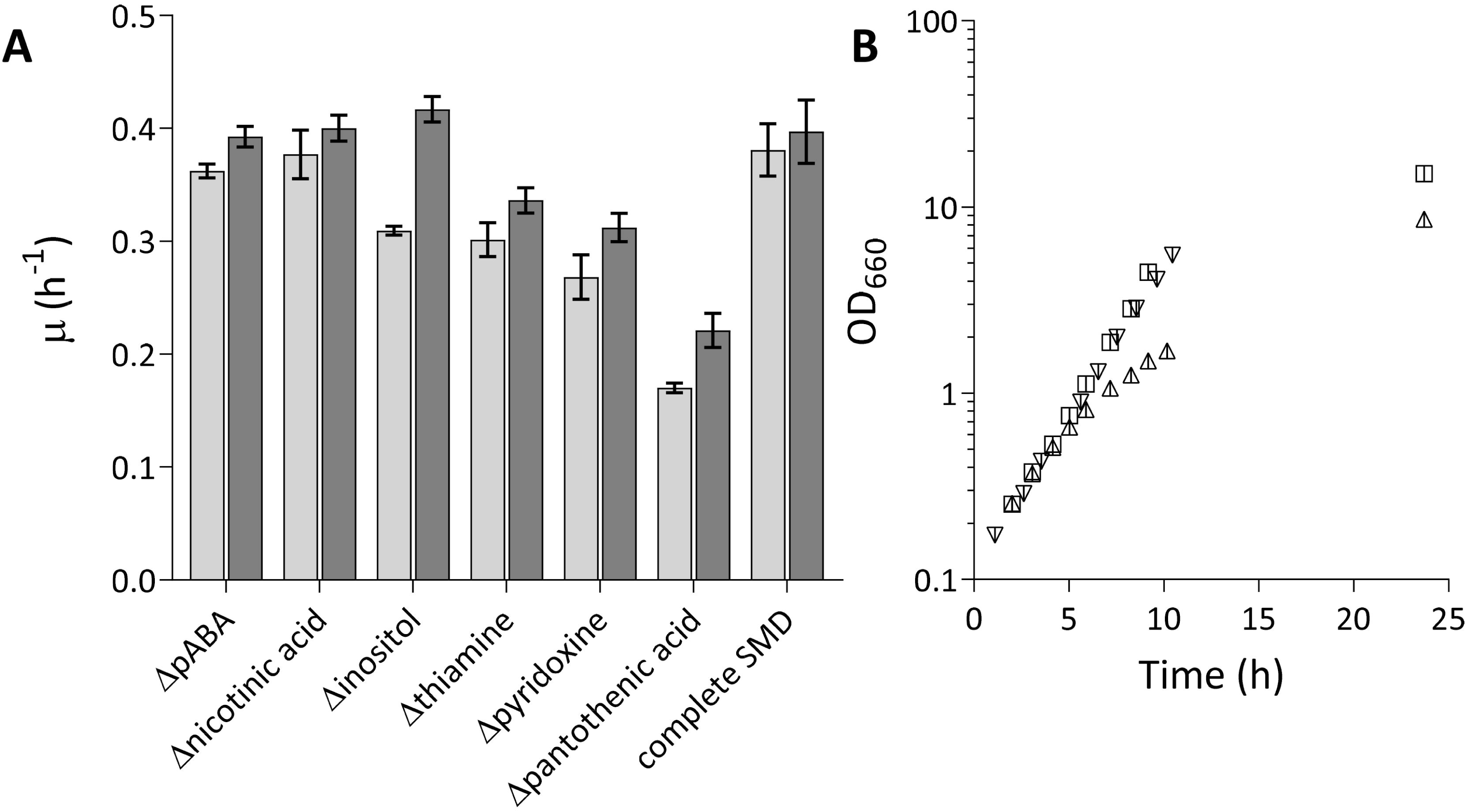
Specific growth rates of *S. cerevisiae* CEN.PK113-7D in aerobic batch cultures on complete SMD and on SMD lacking single vitamins. Growth rate measurements were performed after 3 and 5 consecutive transfers in the same medium. (A). Growth curve of CEN.PK113-7D in complete SMD and at transfer 1 and 3 in SMD lacking *para*-aminobenzoic acid (*p*ABA). In the latter medium a lower specific growth rate was observed at transfer 1 but upon the third transfer, the growth rate was the same as in complete SMD (B). Error bars represent the standard deviation (n = 9 for complete SMD. n = 3 for all other media).

Consistent with the presence in its genome of genes predicted to encode all enzymes involved in biosynthetic pathways for all seven vitamins (Fig. 1, (5)), strain CEN.PK113-7D grew on all SMDΔ versions. On complete SMD, a specific growth rate of 0.38 ± 0.02 h^-1^ was observed, while specific growth rates on SMDΔ lacking biotin, pantothenate, pyridoxine, thiamine or inositol were 95%, 57%, 32%, 22% and 19% lower, respectively. After three transfers, specific growth rates on SMDΔ lacking *p*ABA or nicotinic acid did not differ significantly from the specific growth rate on complete SMD (Fig. 2A). However, in SMDΔ lacking *p*ABA, growth in the first transfer was slower than in the first transfer on complete SMD (Fig. 2B). Extending the number of transfers to five, which corresponded to approximately 33 generations of selective growth, led to higher specific growth rates on several SMDΔ versions (Fig. 2A), suggesting that serial transfer selected for spontaneous faster-growing mutants.

### Adaptive laboratory evolution of CEN.PK113-7D for fast growth in the absence of single vitamins

Serial transfer in independent triplicate aerobic shake-flask cultures on each SMDΔ version was used to select mutants that grew fast in the absence of individual vitamins. Specific growth rates of evolving populations were measured after 5, 10, 23, 38 and 50 transfers and compared to the specific growth rate of strain CEN.PK113-7D grown in complete SMD.

ALE experiments were stopped once the population reached a specific growth rate equal to or higher than 0.35 h^-1^, which represents 90-95% of the specific growth rate of strain CEN.PK113-7D on complete SMD (Fig. 2A) (18–21). As already indicated by the specific growth rates observed after 3 and 5 transfers in SMDΔ (Fig. 2A), few transfers were required for reaching this target in SMDΔ lacking inositol, nicotinic acid or *p*ABA. Conversely, over 330 generations of selective growth were required to reach a specific growth rate of 0.35 h^-1^ on SMDΔ lacking either pantothenic acid, pyridoxine or thiamine (Fig. 3A). At least two single-cell lines were isolated from each of the three independent ALE experiments on each SMDΔ version and the fastest growing single-cell line from each experiment was selected (strains IMS0724-6 from SMDΔ lacking nicotinic acid; IMS0727-9 from SMDΔ lacking *p*ABA; IMS0730-2 from SMDΔ lacking inositol; IMS0733-5 from SMDΔ lacking pantothenate; IMS0736-8 from SMDΔ lacking pyridoxine and IMS0747-9 from SMDΔ lacking thiamine. The specific growth rates of isolates that had been independently evolved in each SMDΔ version did not differ by more than 6%. The largest difference (5.3%) was observed for isolates IMS0733-5 evolved on SMDΔ lacking pantothenate(Fig. 3B).

**Fig. 3:**
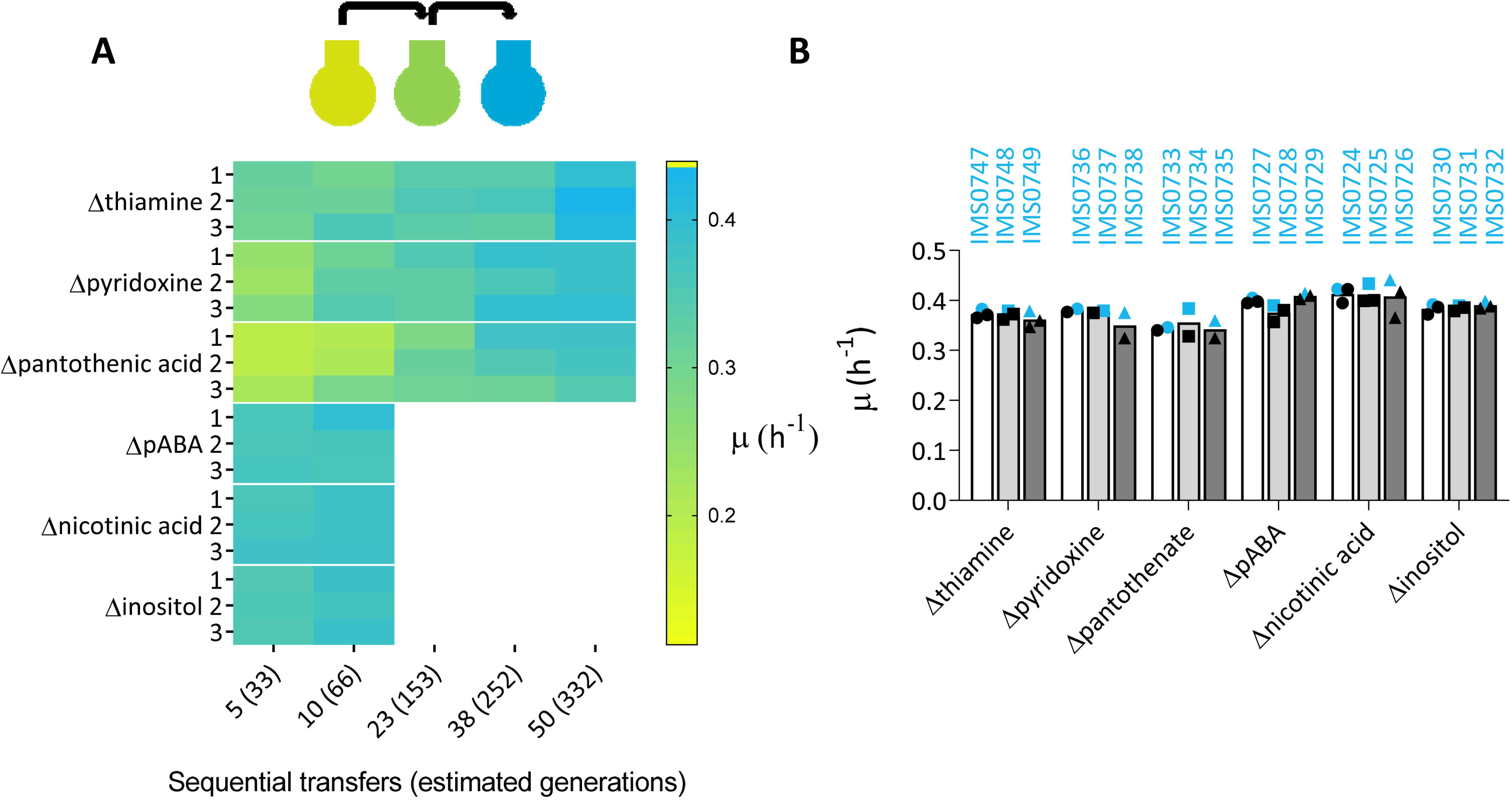
Heat-map showing specific growth rates during ALE of *S. cerevisiae* CEN.PK113-7D on SMD lacking single vitamins. Aerobic serial-transfer experiments on each medium composition were performed in triplicate (rows). The specific growth rate of each evolving population was measured after a specific numbers of sequential transfers (columns). Yellow colour indicates slow growth while cyan indicates a specific growth rate statistically undistinguishable from the positive control (strain CEN.PK113-7D grown on SMD medium with all vitamins) (A). Specific growth rates of single colony isolates from each independent biological replicate evolution line. The fastest-growing isolates, whose genomes were resequenced, are indicated in blue (B).

### Whole-genome sequencing of evolved strains and targets identification

To identify mutations contributing to vitamin independence, the genomes of the sets of three independently evolved isolates for each SMDΔ version were sequenced with Illumina short-read technology. After aligning reads to the reference CEN.PK113-7D genome sequence (22), mapped data were analyzed for the presence of copy number variations (CNV) and single nucleotide variations (SNVs) that occurred in annotated coding sequences.

A segmental amplification of 34 kb (from nucleotide 802500 to 837000) on chromosome VII, which harbours *THI4*, was observed in strain IMS0749 (Fig. 4A) which had been evolved in SMDΔ lacking thiamine. *THI4* encodes a thiazole synthase, a suicide enzyme that can only perform a single catalytic turnover (23). Segmental amplifications on chromosomes III and VIII were observed in strain IMS0725, which had been evolved in SMDΔ lacking nicotinic acid (Fig. 4B.). Since these regions are known to be prone to recombination in the parental strain CEN.PK113-7D (22, 24), their amplification is not necessarily related to nicotinic acid independence.

**Fig. 4:**
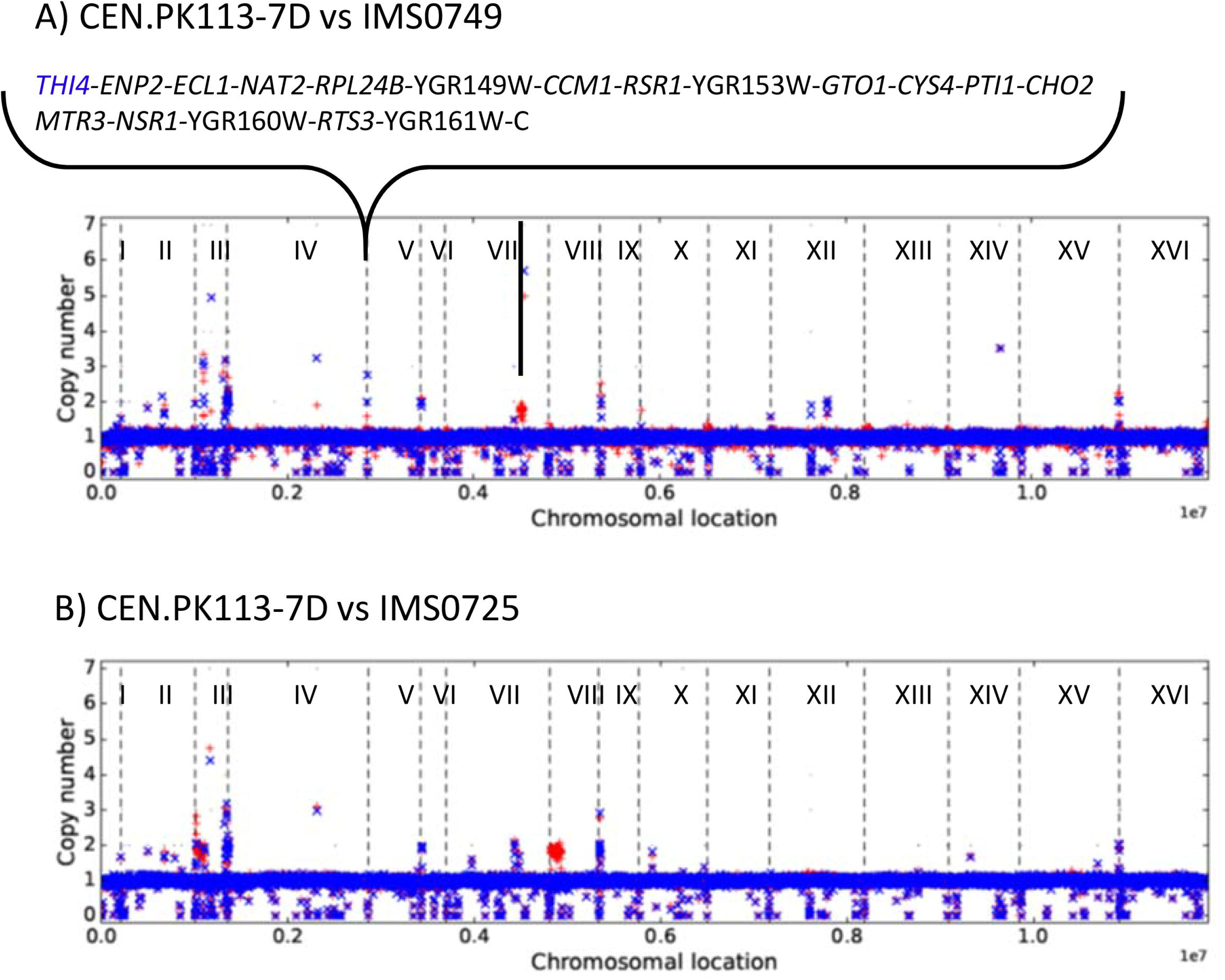
Read coverages across the chromosomes of evolved isolates IMS0725 evolved for nicotinic acid prototrophy (A) and IMS0749 evolved for thiamine prototrophy (B) (in red) compared to read coverage across the chromosomes of CEN.PK113-7D (in blue). Annotated genes found in the amplified region of IMS0749 are indicated.

SNV analysis was systematically performed and data from the three sequenced isolates were compared. To eliminate false positives caused by mapping artifacts, reads of the CEN.PK113-7D strains were mapped back on its own reference assembly. Identified SNVs found were systematically subtracted. SNV analysis was restricted to non-synonymous mutations in coding sequences (Table 2).

**Table 1:** Specific growth rates of best performing single colony isolates obtained from serial-transfer evolution experiments with S. cerevisiae CEN.PK113-7D on SMD and on SMD variants lacking individual B-vitamins. Percentage improvement over the specific growth rate of the parental strain after three transfers in the same medium is also shown (n=1 for each strain).

| *Strain ID* | *Evolution condition* | *Evolution replicate* | *Growth rate ($h^{-1}$)* | *% improvement* |
| --- | --- | --- | --- | --- |
| *IMS0721* | Complete SMD | 1 | 0.443 | 17 |
| *IMS0722* | Complete SMD | 2 | 0.423 | 11 |
| *IMS0723* | Complete SMD | 3 | 0.419 | 10 |
| *IMS0747* | No thiamine | 1 | 0.383 | 35 |
| *IMS0748* | No thiamine | 2 | 0.379 | 30 |
| *IMS0749* | No thiamine | 3 | 0.379 | 38 |
| *IMS0736* | No pyridoxine | 1 | 0.383 | 45 |
| *IMS0737* | No pyridoxine | 2 | 0.379 | 44 |
| *IMS0738* | No pyridoxine | 3 | 0.376 | 48 |
| *IMS0733* | No pantothenate | 1 | 0.346 | 149 |
| *IMS0734* | No pantothenate | 2 | 0.384 | 155 |
| *IMS0735* | No pantothenate | 3 | 0.359 | 159 |
| *IMS0724* | No nicotinic acid | 1 | 0.423 | 4 |
| *IMS0725* | No nicotinic acid | 2 | 0.434 | 2 |
| *IMS0726* | No nicotinic acid | 3 | 0.441 | 2 |
| *IMS0730* | No inositol | 1 | 0.392 | 12 |
| *IMS0731* | No inositol | 2 | 0.389 | 24 |
| *IMS0732* | No inositol | 3 | 0.399 | 16 |
| *IMS0727* | No pABA | 1 | 0.405 | 6 |
| *IMS0728* | No pABA | 2 | 0.389 | 5 |
| *IMS0729* | No pABA | 3 | 0.414 | 4 |

**Table 2:**
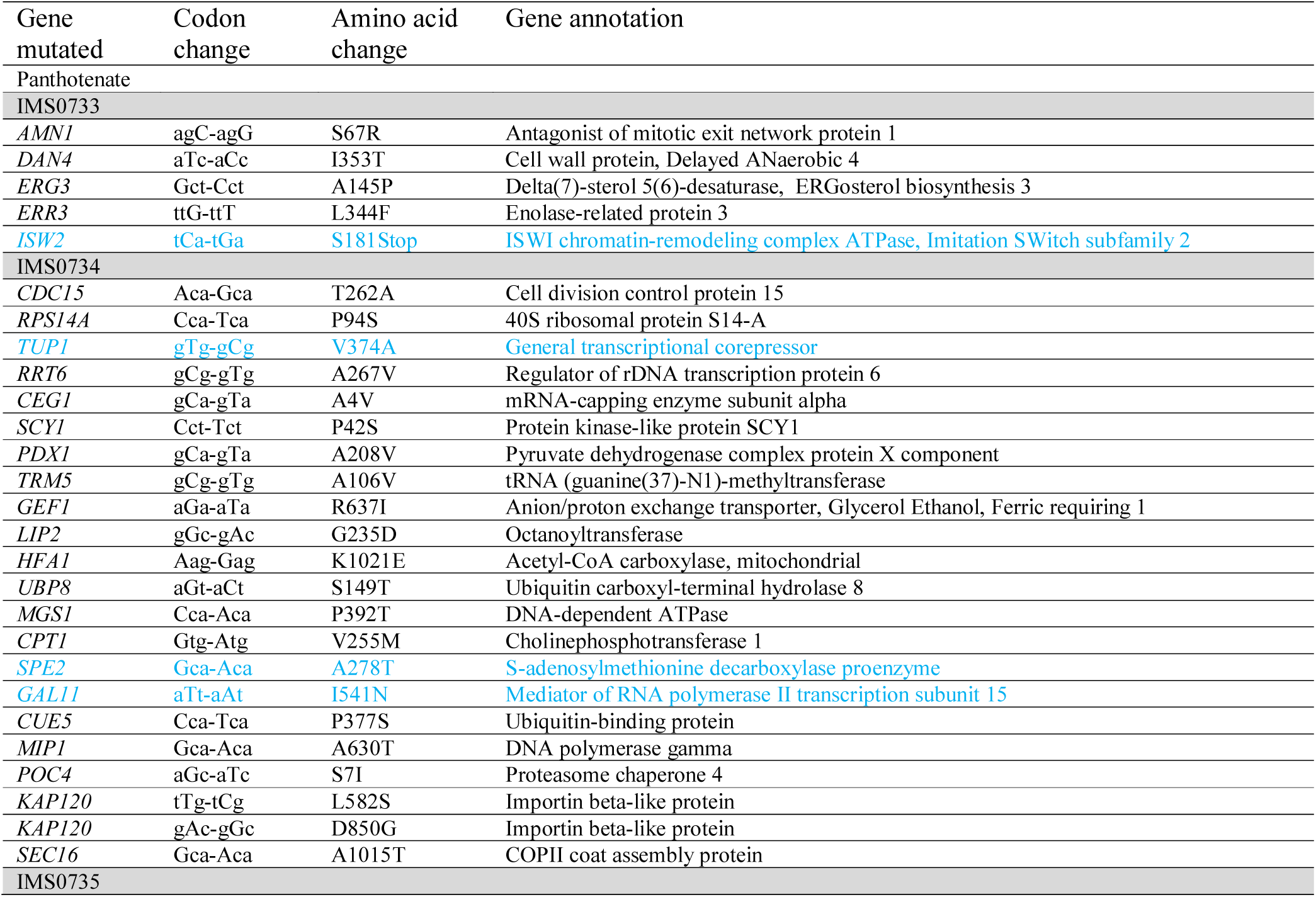
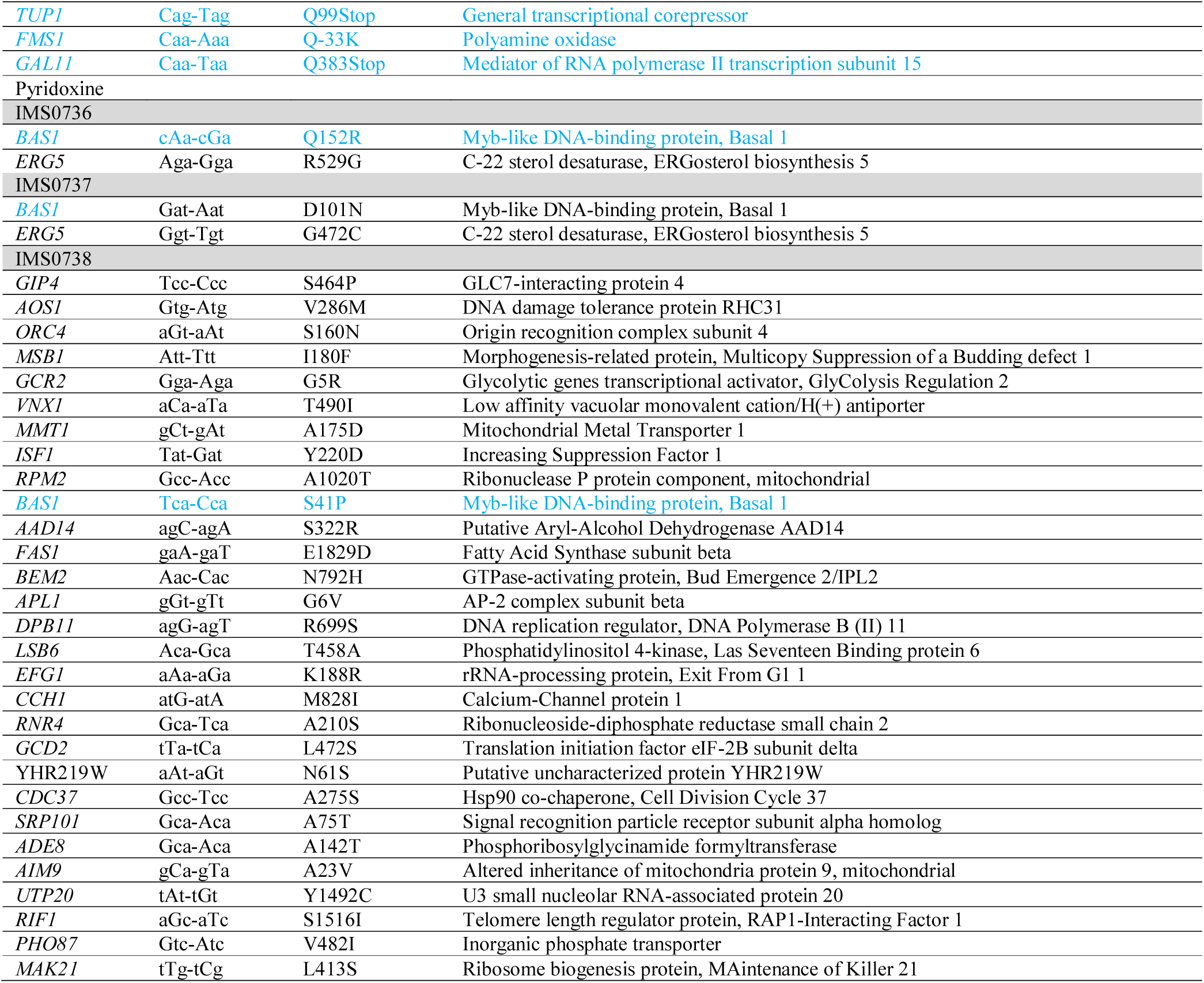
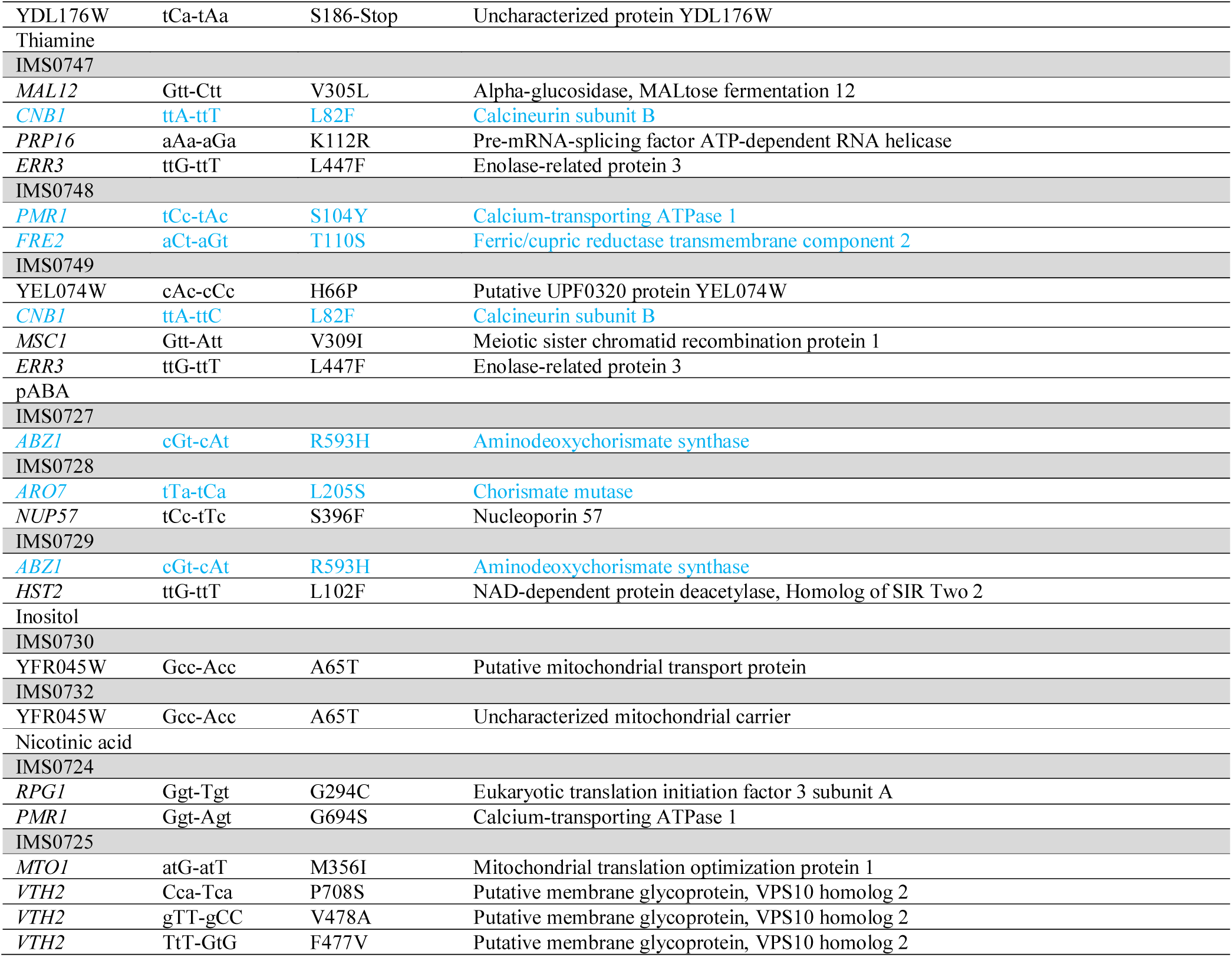
Non-conservative mutations found in single colony isolates obtained from serial-transfer evolution experiments with S. cerevisiae CEN.PK113-7D on on SMD variants lacking individual B-vitamins. Mutations that were chosen for subsequent reverse engineering experiments are shown in blue. S. cerevisiae strains IMS0731 and IMS0726 evolved for fast myo-inositol and nicotinic acid-independent growth, respectively, did not reveal non-conservative mutations and were not included in the table.

In three out of the six isolates from ALE experiments in SMDΔ lacking nicotinic acid or inositol, no non-synonymous SNVs were detected (Fig. 5). One strain (IMS0724) from a serial transfer experiment on SMDΔ lacking nicotinic acid showed SNVs in *RPG1* and *PMR1*, while a second strain (IMS0725) showed SNVs in *MTO1* and *VTH2*. A mutation in YFR054W was identified in a single strain (IMS0730) evolved for inositol-independent growth. The absence of mutations in several strains subjected to serial transfer in SMDΔ lacking nicotinic acid or inositol, is consistent with the fast growth of the parental strain CEN.PK113-7D in these media (Fig. 2A).

**Fig. 5:**
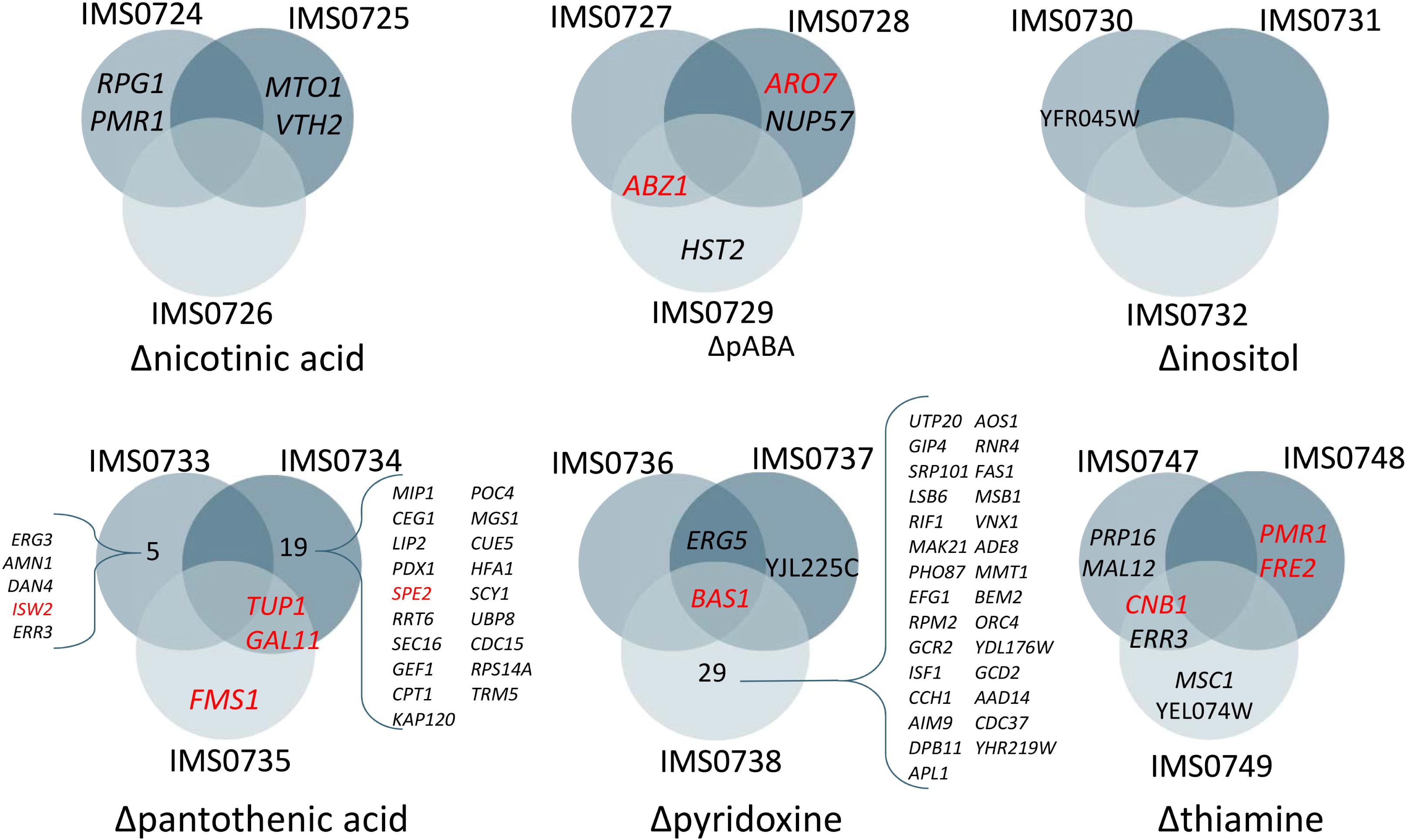
Venn diagrams showing non-synonymous mutations found in coding regions of isolated strains from the different evolution experiments. Each evolution experiment was performed in triplicate. The Venn diagrams show genes that acquired mutations in multiple independent evolution experiments for a specific medium as well as genes that were affected in a single replicate. Apparent mutations also found in the genome of the parent strain CEN.PK113-7D were subtracted and not shown.

Sequencing of the three isolates evolved in SMDΔ lacking *p*ABA revealed only five SNVs, of which two were in *ABZ1* (strains IMS0727 and 0729) and one in *ARO7* (IMS0728), while SNVs in *NUP57* and *HTS2* were found in strains IMS0728 and IMS0729 respectively (Fig. 5). *NUP57* and *HTS2* could not be directly linked to *p*ABA metabolism. Conversely, Abz1 is an aminodeoxychorismate synthase that directs chorismate towards *p*ABA synthesis and Aro7 is a chorismate mutase that catalyses the first committed reaction towards phenylalanine and tyrosine and thereby diverts chorismate from *p*ABA synthesis (Fig.1)(25, 26). These two SNVs therefore represented clear targets for reverse engineering.

In line with the much longer ALE experiments (approximately 332 generations), strains evolved in SMDΔ lacking thiamine, pantothenate or pyridoxine showed larger numbers of SNVs, with a maximum number of 30 SNVs in the isolate IMS0738 evolved SMDΔ lacking pyridoxine (Table 2 and Fig. 5).

Evolution on SMDΔ lacking thiamine did not yield mutations that affected the same gene in all three independently evolved isolates. However, strains IMS0747 and IMS0749 shared SNVs in *CNB1* and *ERR3*. A third isolate, strain IMS0748, contained two SNVs in *PMR1* and *FRE2*. *CNB1*, *PMR1* and *FRE2* all encode proteins that have been implicated in divalent cation homeostasis (27–31).

Isolates IMS0736 and IMS0737, which had been evolved in SMDΔ lacking pyridoxine harboured only two and three mutations, respectively, while strain IMS0738 harboured 30 mutations. All three strains carried different mutated alleles of *BAS1*, which encodes a transcription factor involved in regulation of histidine and purine biosynthesis (32, 33). IMS0736 harboured a non-synonymous mutation causing an amino acid change position 152 (Q152R), while SNVs in strains IMS0737 and IMS0738 affected amino acids 101 (D101N) and 41 (S41P), respectively. Based on these results, *BAS1* was identified as priority target for reverse engineering.

Isolates IMS0733 and IMS0735, evolved on SMDΔ lacking pantothenic acid, carried three and five SNVs, respectively, while isolate IMS0734 carried 21 mutations. Isolates IMS0734 and IMS0735 both carried mutations in *TUP1* and *GAL11*, resulting in different single-amino acid changes (Tup1^V374A^ Gal11^I541N^ and Tup1^Q99stop^ Gal11^Q383stop^, respectively). *TUP1* codes for a general transcriptional corepressor (34) while *GAL11* codes for a subunit of the tail of the mediator complex that regulates activity of RNA polymerase II (35). One of the mutations in strain IMS0733 affected *ISW2*, which encodes a subunit of the chromatin remodeling complex (36). These three genes involved in regulatory processes were selected for reverse engineering, along with *SPE2* and *FMS1*. The latter two genes, encoding S-adenosylmethionine decarboxylase (37) and polyamine oxidase (38), are directly involved in pantothenate biosynthesis and were found to be mutated in isolates IMS0734 and IMS0735, respectively.

In summary, based on mutations in the same gene in independently evolved isolates and/or existing information on involvement of affected genes in vitamin biosynthesis, mutations in twelve genese were selected for reconstruction in the parental strain CEN.PK113-7D. These were mutated alleles of *ISW2*, *GAL11*, *TUP1*, *FMS1* and *SPE2* for panthotenate, in *BAS1* for pyridoxine, mutations in *CNB1, PMR1* and *FRE2* as well as overexpression of *THI4* for thiamine and mutations in *ABZ1* and *ARO7* for pABA. Since serial transfer on SMDΔ lacking nicotininic acid or inositol did not consistently yield mutations and the parental strain CEN.PK113-7D already grew fast on these media, no reverse engineering of observed mutations was observed in isolates from those experimens.

### Reverse engineering of target genes mutations and overexpression

To investigate whether the selected targets contributed to the phenotypes of the evolved strains, single point mutations or single-gene overexpression cassettes were introduced in a non-evolved reference strain, followed by analysis of specific growth rate in the relevant SMDΔ variant. For most target genes, a two-step strategy was adopted, so that a single-gene knock-out mutant was constructed in the process (Fig. 6AB). For the *SPE2* mutant strains IMX2308 and IMX2289, point mutations were introduced in a single step (Fig. 6C). The *THI4-*overexpressing strains IMX2290 and IMX2291 were constructed by integrating the overexpression cassette at the YPRcTau3 locus (39) (Fig. 6D). Subsequently, multiple mutations that were found in strains evolved in the same SMDΔ version were combined into single engineered strains to test for additive or synergistic effects.

**Fig. 6:**
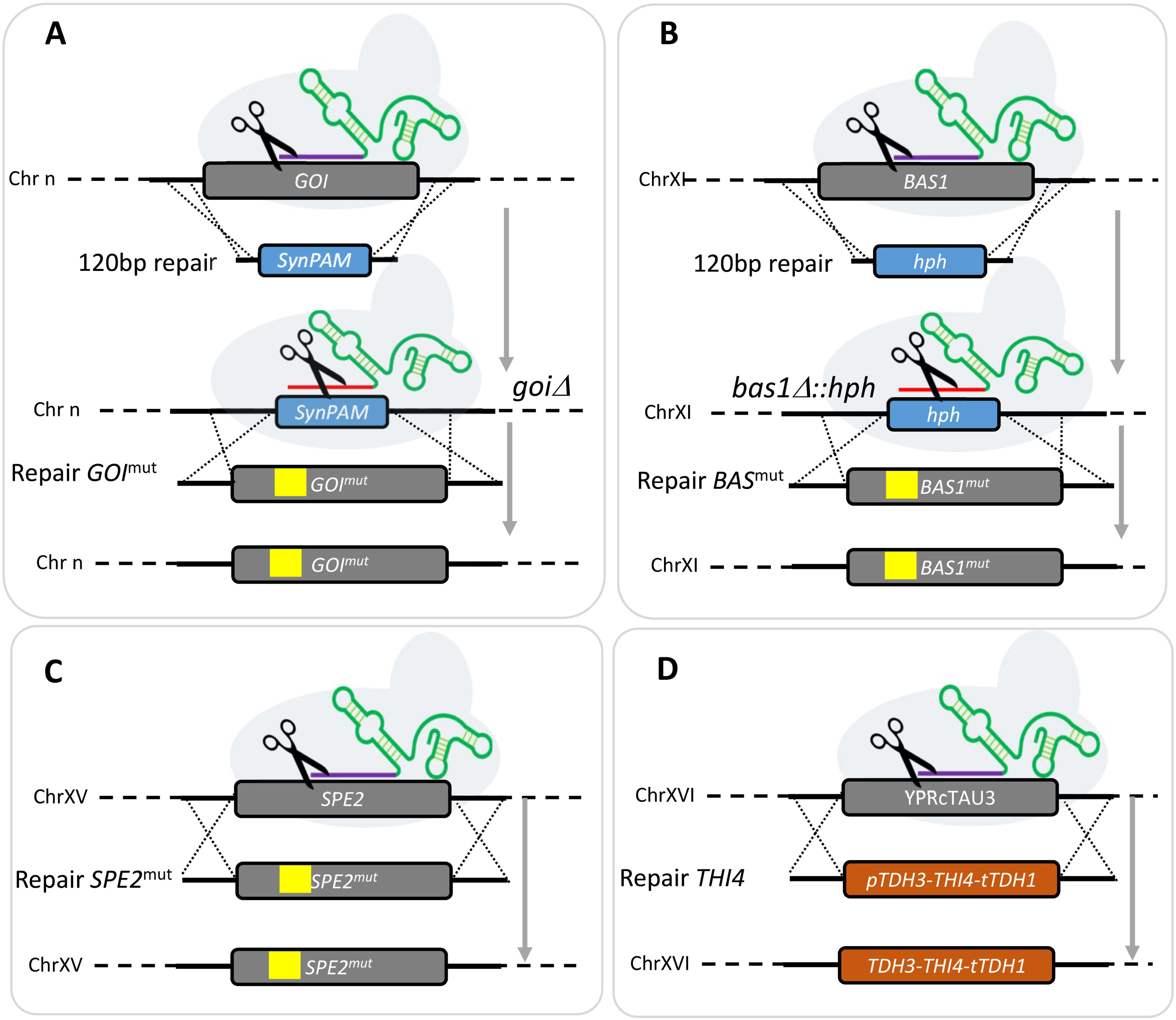
Strain construction strategy for reverse engineering. Most of the single mutation strains were generated in two steps. First the gene of interest (GOI) was replaced by a synthetic 20 bp target sequence and 3 bp PAM sequence (SynPAM). In a second step, the SynPAM was targeted by Cas9 and substituted with the GOI mutant allele (A). The SynPAM approach was not successful in targeting *BAS1.* For this reason, the *BAS1* mutant strains (IMX2135-7) were constructed by first knocking out the gene by replacing it with the antibiotic marker hphNT1 that confers resistance to hygromycin. Then, in a second step, the selection marker was targeted with Cas9 and substituted with a *BAS1* mutant allele (**B**). In the case of *SPE2* mutant strains (IMX2289 and IMX2308), the mutant allele was swapped with the WT allele in a single step (**C**). The *THI4* overexpressing strains IMX2290 and IMX2291 were constructed by integrating a *THI4* overexpression cassette at the YPRcTau3 *locus* (**D**). SNVs are represented by yellow boxes.

### Thiamine

The specific growth rate of *S. cerevisiae* CEN.PK113-7D was only 27% lower in SMDΔ lacking thiamin than in complete SMD (Fig. 2A and 6A). Nevertheless, it took over 300 generations of selective growth to obtain evolved isolates that compensated for this difference (Fig.3 and 6A). The role of mutations in *CNB1*, *FRE2* and *PMR1* in this evolved phenotype was first investigated in the single knock-out strains IMX1721, IMX1722 and IMX1723, respectively. While deletion of *PMR1* deletion negatively affected specific growth rate on SMDΔ lacking thiamine, deletion of either *CNB1* or *FRE2* resulted in an 17% increase of the specific growth rate on this medium relative to CEN.PK113-7D. However, strains IMX1721 (*cnb1*Δ) and IMX1722 (*fre2*Δ) still grew significantly slower than the evolved isolates (Fig.6). Subsequently, the mutated alleles found in the evolved isolates were introduced at the native chromosomal locus, resulting in strains IMX1985 (*CNB1*^L82F)^), IMX1986 (*PMR1*^S104Y^) and IMX1987 (*FRE2*^T110S^). In addition, *THI4* was overexpressed (strain IMX2290) to simulate the copy number increase observed in IMS0749. Strains IMX1987 (*FRE2*^T110S^) and IMX2290 (*THI4*↑) grew as fast as the evolved isolates on SMDΔ lacking thiamine (0.35-0.36h^-1^; Fig. 5A). Combination of these mutated alleles of *PMR1* and *FRE2*, which occurred together in isolate IMS0748, as well as of the two mutations resulting in growth improvement (*FRE2*^T110S^ and *THI4*↑) was also tested. None of these combinations yielded a higher specific growth rate than observed in the evolved strains and in the reversed engineered *FRE2*^T110S^ and *THI4*↑ strains.

### *para*-Aminobenzoic acid

In SMDΔ lacking *p*ABA, strain CEN.PK113-7D grew 50% slower than in complete SMD. However, it took only a few transfers to achieve fast *p*ABA-independent growth. The independently evolved isolates IMX2057 and IMX1989 harboured mutations affecting genes that encode chorismate-utilizing enzymes, the precursor of *p*ABA (*ABZ1*^N593H^ and *ARO7*^L205S^, respectively; Fig. 1). As these strains were able to grow in SMD without amino acid supplementation, these mutations affecting did not cause a complete loss of function.

However, they might well affect distribution of chorismate over *p*ABA and aromatic-amino-acid biosynthesis (25, 26). Introduction of either *ABZ1*^N593H^ or *ARO7*^L205S^, while replacing the corresponding wild-type allele, eliminated the slower growth observed in strain CEN.PK113-7D during the first transfer on SMDΔ lacking *p*ABA. Specific growth rates of these reverse engineered strains IMX2057 (*ABZ1*^R593H^) and IMX1989 (*ARO7*^L205S^) were not statistically different from those of the corresponding evolved isolates (Fig. 6B).

### Pantothenic acid

Omission of pantothenic acid from SMD led to a 57% lower specific growth rate of strain CEN.PK113-7D than observed in complete SMD (Fig. 2 and Fig 6C). Out of a total number of 29 mutations found in three independently evolved isolates that showed fast growth in SMDΔ lacking pantothenate, SNVs in *ISW2*, *GAL11*, *TUP1*, *SPE2*, and *FMS1* were analysed by reverse engineering. Single deletion of *SPE2*, *FMS1* and *GAL11* resulted in an inability to grow on SMDΔ lacking pantothenate. This result was anticipated for the *spe2*Δ and *fms1*Δ mutants, in view of the roles of these genes in pantothenate biosynthesis. However, *GAL11* has not previously been implicated in pantothenate biosynthesis. The *gal11*Δ strain was conditional as the mutant did grow on complex YPD and SMD media. Of the remaining two deletion mutants, the *tup1*Δ strain IMX1817 showed a 68 % higher specific growth rate on SMDΔ than strain CEN.PK113-7D (Fig. 6C), while deletion of *ISW2* did not result in faster growth on this medium (Fig. 6C). Of seven SNVs that were individually expressed in the non-evolved strain background, only the *GAL11* ^Q383Stop^ mutation found in IMS0735 supported a specific growth rate of 0.33 h^-1^ on SMDΔ lacking pantothenate that was only 8% lower to that of the evolved isolates.

Combination of the *GAL11*^Q383Stop^ mutation with *TUP1*^V374A^, *TUP1*^Q99Stop^, and *SPE2* or *TUP1* with *FMS1* did not lead to additional improvement, indicating that the *GAL11*^N383Stop^ mutation was predominantly responsible for the improved growth of evolved strains IMS0734 and IMS0735 in the absence of pantothenate.

### Pyridoxine

Strain CEN.PK113-7D grew 35% slower on SMDΔ lacking pyridoxine than on complete SMD. Three different mutated alleles of *BAS1* were identified in strains that had been independently evolved for fast growth on the former medium (Table 2). Deletion, in a non-evolved reference strain, of *BAS1* (IMX2128) did not result in faster pyridoxine-independent growth (Fig. 6D). Individual expression of the evolved *BAS1* alleles in strain IMX2128 yielded strains IMX2135 (*BAS1*^Q152R^), IMX2136 (*BAS1*^D101N^) and IMX2137 (*BAS1*^S41P^). All three *BAS1* mutant strains grew faster on SMDΔ lacking pyridoxine than strain CEN.PK113-7D, reaching specific growth rates on this medium that were not significantly different from the average of those of evolved strains IMS736, IMS737, and IMS738 (Figure 5D). These results suggest that *BAS1*, which has previously been shown to be involved in regulation of purine and histidine biosynthesis (32, 33), may also be involved in regulation of pyridoxine biosynthesis in *S. cerevisiae*.

## Discussion

### Vitamin requirements of *S. cerevisiae*

Most *S. cerevisiae* genomes harbor the full complement of genes required for synthesis of the seven B-vitamins that are commonly included in chemically defined media for yeast cultivation (CDMY, for a recent review see (5, 10). Previous studies indicated that presence of a complete set of biotin biosynthesis genes supported only slow growth on CDMY. The present study shows that, similarly, none of the other six B-vitamins included in CDMY (inositol, nicotinic acid, pantothenic acid, *p*ABA, pyridoxine and thiamine) are strictly required for growth. Remarkably, the impact of individually eliminating these six vitamins from a glucose-containing CDMY differently affected specific growth rates in aerobic, glucose-grown cultures, with growth-rate reductions varying from 0 to 57 %. It should, however, be noted that requirements for these growth factors, which for aerobic yeast cultivation cannot be formally defined as vitamins and that their absolute and relative requirements may will be condition- and strain dependent. For example, it is well documented that synthesis of nicotinic acid by *S. cerevisiae* is strictly oxygen dependent (40). The dataset compiled in the present study will, hopefully, serve as reference for investigating vitamin requirements of diverse natural isolates, laboratory and industrial strains and thereby help to obtain a deeper understanding of the genetics and ecology of vitamin prototrophy and vitamin biosynthesis in *S. cerevisiae*.

### ALE and reverse engineering for identifying genes involved in fast B-vitamin indepent growth

A serial transfer strategy was applied to select for spontaneous mutants that grew as fast in aerobic batch cultures on CDMY lacking either inositol, nicotinic acid, pyridoxine, thiamine, pantothenic acid, or *para*-aminobenzoic acid as in CDMY containing all these six vitamins as well as biotin. In the ALE experiments on media lacking nicotinic acid or inositol, fast growth was observed within a few cycles of batch cultivation and not all fast-growing strains were found to contain mutations. These observations indicated that, under the experimental conditions, the native metabolic and regulatory network of *S. cerevisiae* was able to meet cellular requirements for fast growth in the absence of these ‘vitamins’.

As demonstrated in other ALE studies, performing independent replicate evolution experiments helped in identifying biologically relevant mutations upon subsequent whole-genome sequencing (11, 12). The power of this approach is illustrated by the ALE experiments that selected for pyridoxine-independent growth, in which the independently evolved mutants IMS0736 and IMS0738 harboured 2 and 30 mutated genes, respectively, of which only *BAS1* also carried a mutation in a third, independently sequenced isolate (Fig. 5).

In total, the role of 12 genes that were found to be mutated in the ALE experiments were selected for further analysis by reverse engineering of the evolved alleles and/or deletion mutations in the parental, non-evolved genetic background (Fig. 5 and Table 2). These genes comprised three groups; i) genes encoding enzymes known or inferred to be involved in the relevant vitamin synthesis pathway (*SPE2* and *FMS1* for pantothenate, *THI4* for thiamine, *ABZ1* and *ARO7* for *p*ABA), ii) genes encoding transcriptional regulator proteins (*TUP1* and *GAL11* for pantothenate and *BAS1* for pyridoxine) and iii) non-transcriptional-regulator proteins whose functions have not previously been associated with vitamin biosynthesis (*ISW2* for pantothenate and *CNB1, PMR1* and *FRE2* for thiamine).

Of the first group of mutations defined above, only those in *SPE2* and *FMS1* were not found to contribute to faster growth in the absence of the relevant vitamin. The second group yielded interesting information on regulation of vitamin biosynthesis in *S. cerevisiae*. In particular, the key role of mutations in *BAS1* in enabling fast pyridoxine-independent growth and the role of *GAL11* mutations in fast pantothenate-independent growth dependency provided interesting insights and leads for further research.

The *S. cerevisiae* transcriptional activator Bas1 is involved in regulation of purine and histidine (32, 33). Interestingly, Bas1 has also is also involved in repression of genes involved in C1 metabolism and of *SNZ1* (41). Snz1 is a subunit of a two-component pyridoxal-5’-phosphate synthase, which catalyses the first step of the synthesis of pyridoxal-5-phosphate, the active form of pyridoxine in *S. cerevisiae* (42). Interrogation of the Yeastract database (43) for occurrence of transcription binding site in promoter regions of pyridoxine-biosynthesis genes confirmed the link already established between *BAS1* and *SNZ1* (41, 44). Moreover, this analysis revealed that all pyridoxine biosynthesis genes in *S. cerevisiae* contain a consensus Bas1 cis-regulatory binding motif (Fig. 8). Consistent with the regulatory role of Bas1 on *SNZ1* expression, Bas1 has been experimentally shown to repress transcription of genes involved in pyridoxine biosynthesis (45). The mutations found in *BAS1* may therefore have attenuated Bas1-mediated repression of pyridoxine biosynthetic genes and, thereby, enabled increased pyridoxine biosynthesis.

**Fig. 8:**
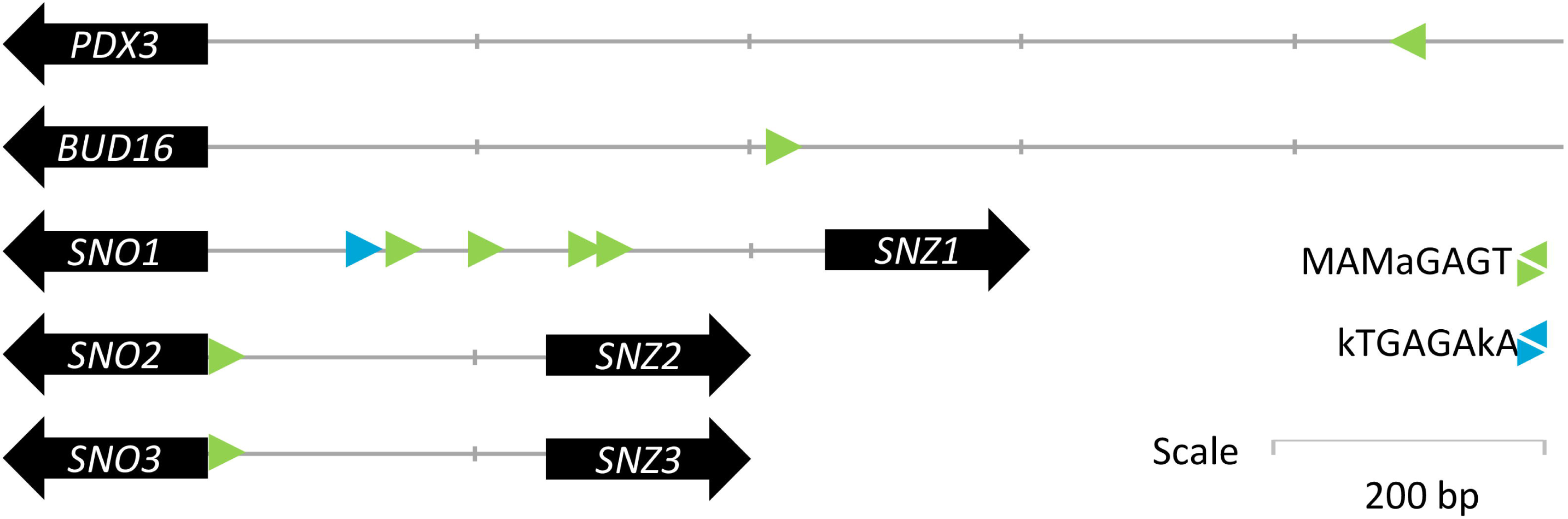
Schematic representation of Bas1 binding sites in promoter regions of genes involved in pyridoxal-5-phosphate biosynthesis. The two Bas1 consensus binding sequences MAMaGAGT and kTGAGAkA (**73**) are shown in green and blue respectively. Scale bar indicates 200 bp.

ALE experiments in pantothenate-free medium yielded different mutations in *TUP1* and *GAL11,* two major components of the yeast regulatory machinery. *TUP1* encodes a general transcriptional repressor that, in a complex with Cyc8, modifies chromatin structure such that genes are repressed (46–48). *GAL11* (also known as *MED15*) encodes a subunit of the mediator complex required for initiation by RNA polymerase-II and consequently plays a critical role in transcription of large RNA polymerase-II dependent genes (49, 50). Despite its involvement in general cellular transcriptional regulation, *GAL11* is not an essential gene for growth in comlete medium (51). The inability of a *gal11*Δ strain to grow on glucose synthetic medium without pantothenate represents the first indication for a possible involvement of Gal11 in regulation of pantothenate metabolism. Gal11 interacts with transcriptional activators through various peptidic segments including an N-terminal KIX domain. This region shows homology with the B-box motif found in the mammalian activating protein SRC-1 and is essential for recruitment of the mediator complex by other regulatory proteins (e.g. Gcn4) (52). Of two mutations found in *GAL11*, the most potent was a nonsense mutation at nucleotide 383. In contrast to a *gal11*Δ strain A reverse engineered strain carrying this premature stop codon grew on SMDΔ pantotenate, which indicates that the *GAL11*^R383Stop^ allele encodes a functional peptide. Such a functional, truncated Gal11 version has not been previously described is sufficiently long to include a complete KIX domain (AA_9_-AA_86_) for recruitment of the RNA polymerase-II machinery by an as yet unidentified transcription factor involved in regulation of pantothenate biosynthesis. Further research is required to resolve and understand the role of the wild-type and evolved alleles of *GAL11* in regulation of pantothenate metabolism.

A third group of non-transcription factor genes had not yet been associated with the biosynthesis of vitamins. Reverse engineering of a mutation in *ISW2*, which encodes an ATP-dependent DNA translocase involved in chromatin remodeling (53) identified in the pantothenate evolution did not yield a growth improvement, but we cannot exclude that this mutation in association with *ERG3, AMN1*, *DAN4,* and *ERR3* identified in IMS0733 (Fig. 5 and 6C) could have a significant impact but systematic combinatorial analysis of the mutations was not performed.

Mutations in *CNB1*, *PMR1* and *FRE2* identified in evolved isolates all improved growth of *S. cerevisiae* in the absence of thiamine (Fig. 7A). These three genes all encode proteins involved in metal homeostasis, Fre2 is a ferric or cupric reductase (54), Cnb1 is the regulatory B-subunit of calcineurin, a Ca^2+^/calmodulin-regulated type 2B protein phosphatase which regulates the nuclear localization of Crz1. This transcription factor influences expression of a large number of genes. Its targets include *PMR1*, which encodes a high-affinity Ca^2+^/Mn^2+^ P-type ATPase involved in Ca^2+^ and Mn^2+^ transport into the Golgi (29, 55). Neither of these three genes have hitherto been directly associated with thiamine. However, thiamine pyrophosphokinase (Thi80), thiamin phosphate synthase (Thi6) and hydroxymethylpyrimidine phosphate (Thi21 and Thi20) all require Mg^2+^ or Mn^2+^ as co-factors (56, 57). At low concentration, Mn^2+^ was shown to be a stronger activator of Thi80 than Mg^2+^(58). In an ALE experiment with engineered xylose-fermenting assimilating *S. cerevisiae*, a non-sense mutation or deletion of *PMR1* caused selectively and strongly increased intracellular concentrations of Mn^2+^, which was the preferred metal ion for the heterologously expressed *Piromyces* xylose isomerase (59). Although intracellular metal ion concentrations were not measured in the current study, the different phenotypes of a *pmr1*Δ deletion strain (IMX1722) and a *PMR1*^S104Y^ strain (IMX1986) (Fig. 7A) indicate that the latter mutation does not act through a massive increase of the intracellular Mn^2+^ concentration.

**Fig. 7:**
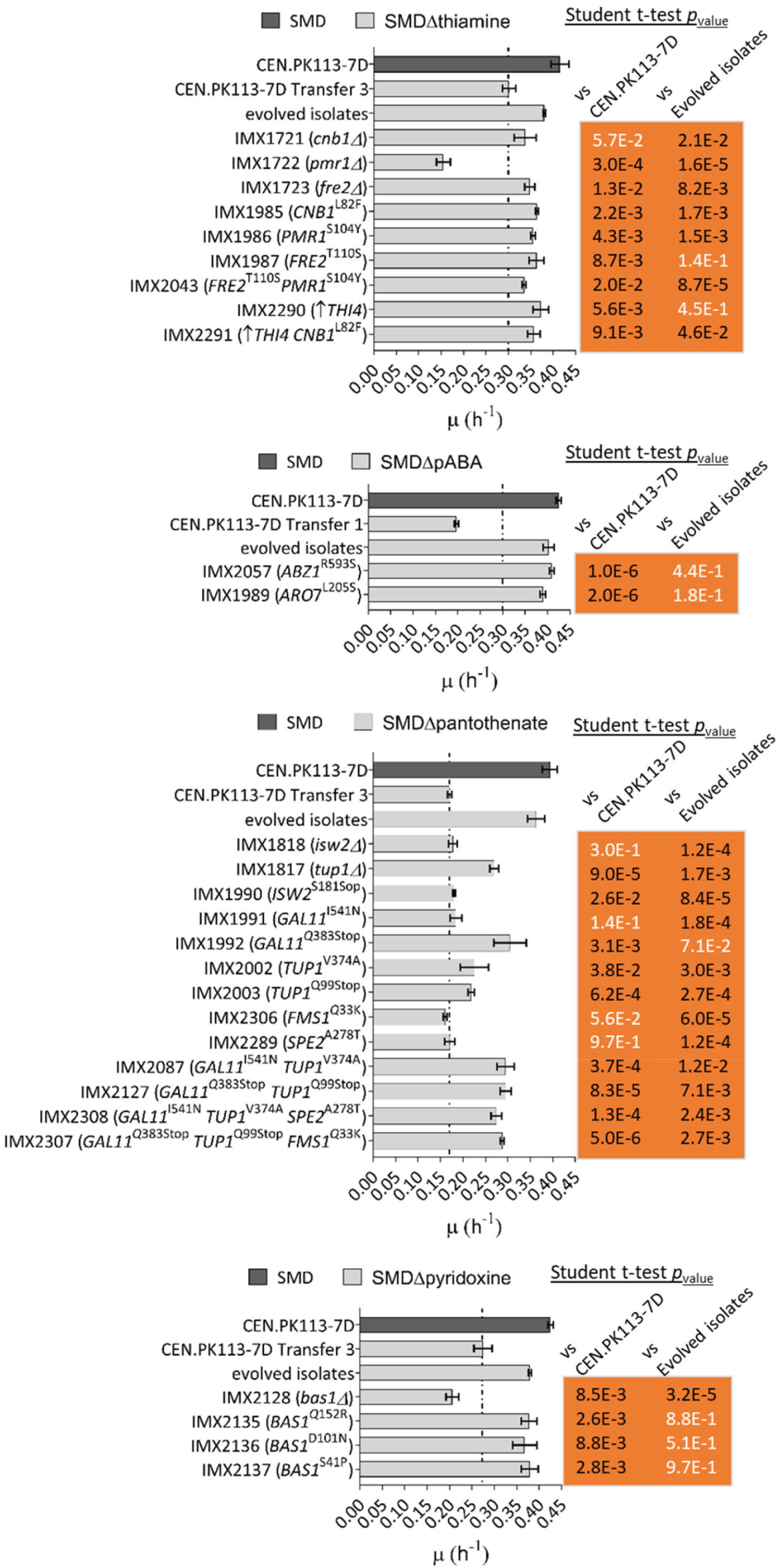
Specific growth rates of engineered *S. cerevisiae* strains carrying one or multiple gene deletions or reverse engineered mutations in SMD media lacking thiamine (A), pABA (B), pantothenic acid (C) and pyridoxine (D). Specific growth rates of *S. cerevisiae* CEN.PK113-7D grown in complete SMD and evolved CEN.PK113-7D in SMD medium lacking the relevant vitamin are shown as references. The specific growth rate of strain CEN.PK113-7D in SMD medium lacking the relevant vitamin is shown and highlighted by a vertical line to help to visualize improved performance of engineered strains. Error bars represent the standard deviation (n = 9 for complete SMD, n = 6 for strain IMX1721, otherwise n=3). A Student t-test was performed to compare the wild-type and evolved CEN.PK113-7D growth rate to the engineered strains growth rate and non-significant differences are indicated with white letters (p-value > 0.05).

In *S. cerevisiae,* synthesis of the thiazole moiety of thiamine biosynthesis involves sulfide transfer from an active-site cysteine (Cys205) residue of the thiazole synthase Thi4. This sulfur transfer reaction is iron-dependent and generates inactive enzyme by formation of a dehydroalanine. Fe(II) plays an essential role in this sulfide transfer, which remains poorly understood (23). Further research is needed to investigate if the *FRE2* mutation in strain IMS0749 in some way increases the efficiency of the reaction catalyzed by the energetically single-turnover enzyme Thi4 and to resolve the role of metal homeostasis in vitamin biosynthesis.

### Towards mineral media for cultivation of S. cerevisiae

With the exception of the carbon and energy source for growth, B-vitamins are the sole organic ingredients in standard CDMY recipes for aerobic cultivation of wild-type and industrial *S. cerevisiae* strains. In view of the chemical instability of some of these compounds, vitamin solutions cannot be autoclaved along with other medium components but are usually filter sterilized. In research laboratories and in particular in industrial processes, the costs, complexity and contamination risks associated with the use of vitamins is significant. Complete elimination of vitamins from CDMY, without compromising on specific growth rate, yield or productivity, could therefore result in consideral cost and time savings as well as in improved standardization and robustness of cultivation procedures.

The present study demonstrates that, by ALE as well as introduction of small sets of defined mutations into *S. cerevisiae*, it is is possible to achieve specific growth rates in single-vitamin depleted CDMY that are close or identical to those found in CDMY supplemented with a complete vitamin mixture. While these results represent a first step towards the construction of completely prototrophic growth of *S. cerevisiae* and related yeasts, further research is required which trade-offs are incurred upon simultaneous introduction of the genetic interventions identified in this study and how they can be mitigated. This issue may be particularly relevant for mutations that affect genes involved in global regulation processes (50, 60), which may interfere with other cellular processes. In addition, simultaneous high-level expression of multiple enzymes with low-catalytic turn-over numbers, with the suicide enzyme Thi4 (23, 61, 62) as an extreme example, may affect cell physiology due to the required resource allocation (63, 64).

In such cases, it may be necessary to expand metabolic engineering strategies beyond the native metabolic and regulatory capabilities of *S. cerevisiae* by expression of heterologous proteins and/or pathways with more favourable characteristics.

## Materials and Methods

### Strains, media and maintenance

The *S. cerevisiae* strains used and constructed in this study are shown in Table 3 and they all derive from the CEN.PK lineage (16, 65). Yeast strains were grown on synthetic medium with ammonium sulfate as a nitrogen source (SM) or YP medium (10 g/L Bacto yeast extract, 20 g/L Bacto peptone) as previously described (2). SM and YP media were autoclaved at 121°C for 20 min. Then, SM medium was supplemented with 1 ml/L of filter-sterilized vitamin solution (0.05 g/L D-(+)-biotin, 1.0 g/L D-calcium pantothenate, 1.0 g/L nicotinic acid, 25 g/L myo-inositol, 1.0 g/L thiamine hydrochloride, 1.0 g/L pyridoxol hydrochloride, 0.20 g/L 4-aminobenzoic acid). Vitamin drop-out media were prepared using vitamin solutions lacking either thiamine, pyridoxine, pantothenic acid, inositol, nicotinic acid or *para*-aminobenzoic acid, yielding SMΔthiamine, SMΔpyridoxine, SMΔpantothenic acid, SMΔinositol, SMΔnicotinic acid, SMΔ*p*ABA respectively. A concentrated glucose solution was autoclaved at 110 °C for 20 min and then added to the SM and YP medium at a final concentration of 20 g/L, yielding SMD and YPD, respectively. 500 ml shake flasks containing 100 ml medium and 100 ml shake flasks containing 20 ml medium were incubated at 30 °C and at 200 rpm in an Innova Incubator (Brunswick Scientific, Edison, NJ). Solid media were prepared by adding 1.5% Bacto agar and, when indicated, 200 mg/L G418 or 200 mg/L hygromycin. *Escherichia coli* strains were grown in LB (10 g/L Bacto tryptone, 5 g/L Bacto yeast extract, 5 g/L NaCl) supplemented with 100 mg/L ampicillin or kanamycin. *S. cerevisiae* and *E. coli* cultures were stored at −80 °C after the addition of 30% v/v glycerol.

**Table 3:**
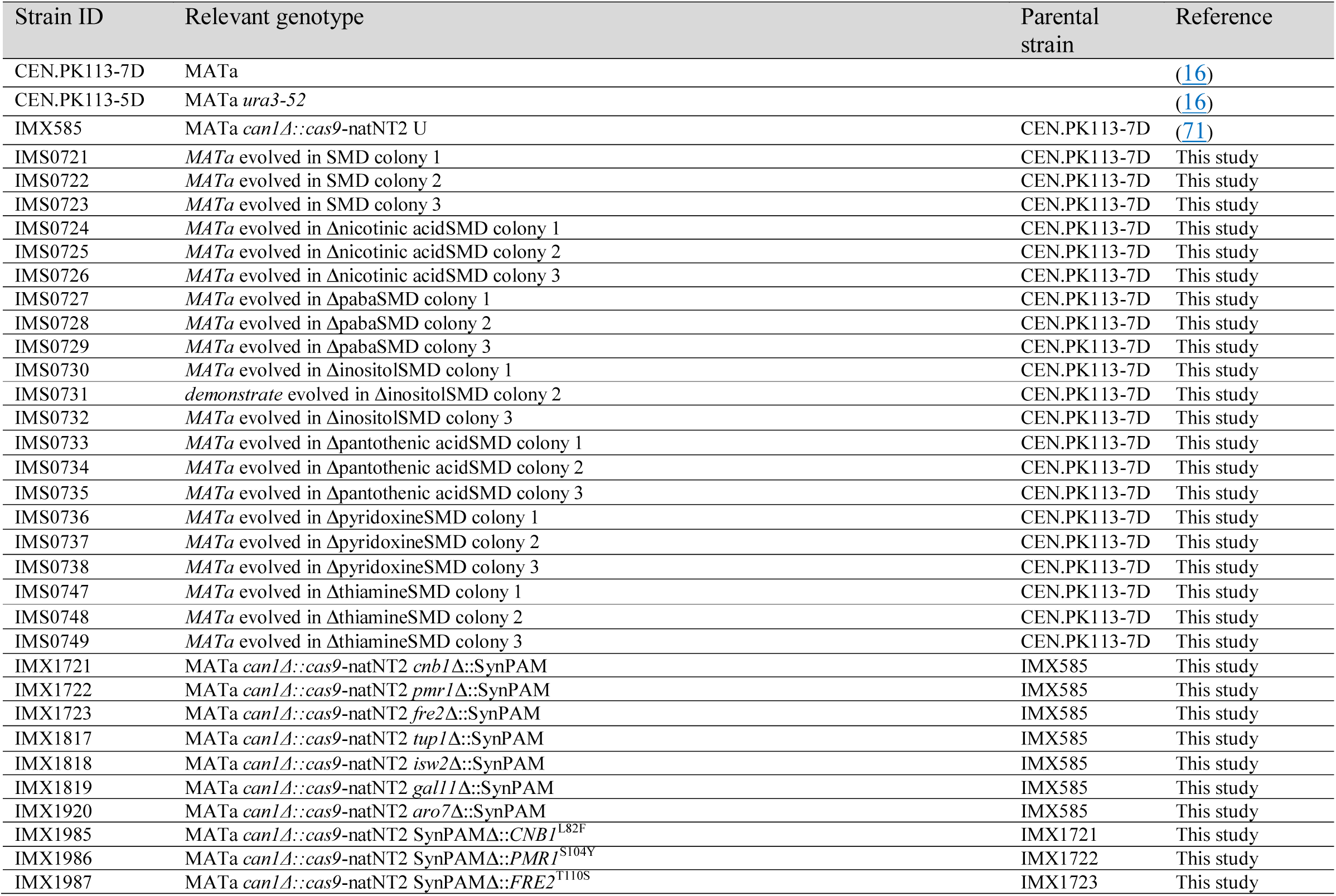
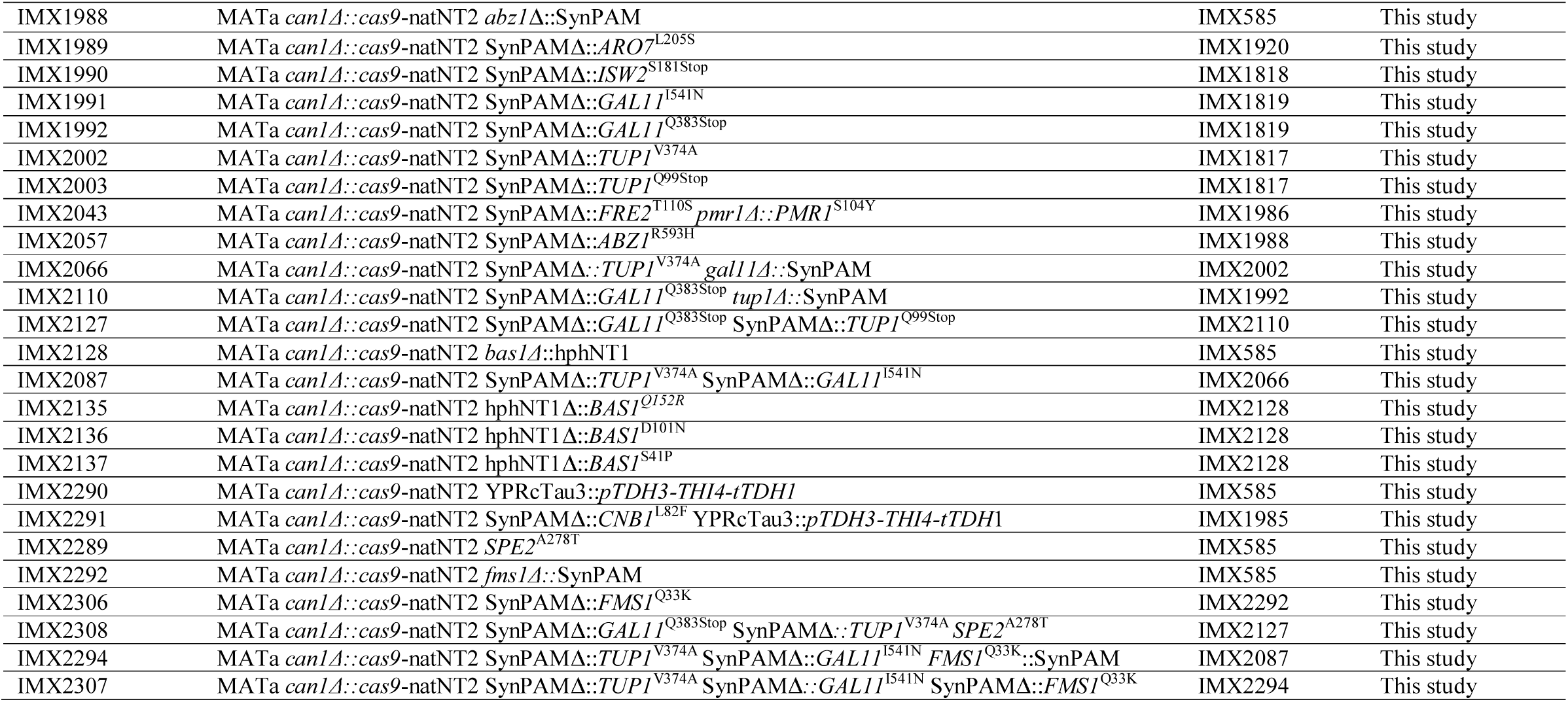
*Saccharomyces cerevisiae* strains used in this study.

### Molecular biology techniques

PCR amplification of DNA fragments with Phusion Hot Start II High Fidelity Polymerase (Thermo Scientific, Waltham, MA) and desalted or PAGE-purified oligonucleotide primers (Sigma-Aldrich, St Louis, MO) was performed according to manufacturers’ instructions. DreamTaq polymerase (Thermo Scientific) was used for diagnostic PCR. Primers used in this study are shown in Table 5. PCR products were separated by gel electrophoresis using 1 % (w/v) agarose gels (Thermo Scientific) in TAE buffer (Thermo Scientific) at 100 V for 25 min and purified with either GenElutePCR Clean-Up Kit (Sigma-Aldrich) or with Zymoclean Gel DNA Recovery Kit (Zymo Research, Irvine, CA). Plasmids were purified from *E. coli* using a Sigma GenElute Plasmid Kit (Sigma Aldrich). Plasmids used in this study are shown in Table 4. Yeast genomic DNA was isolated with the SDS/LiAc protocol (66). Yeast strains were transformed with the lithium acetate method (67). Four to eight single colonies were re-streaked three consecutive times on selective media and diagnostic PCR were performed in order to verify their genotype. *E. coli* XL1-blue was used for chemical transformation (68). Plasmids were then isolated and verified by either restriction analysis or by diagnostic PCR.

**Table 4:**
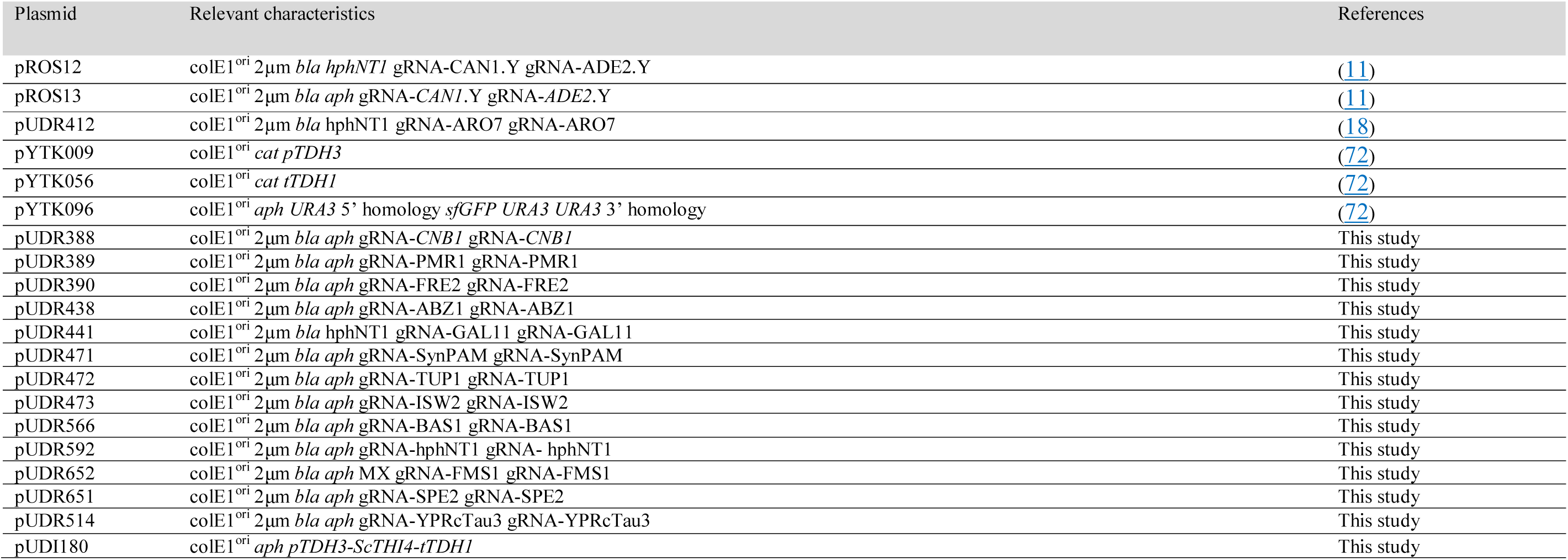
plasmids used in this study.

**Table 5:**
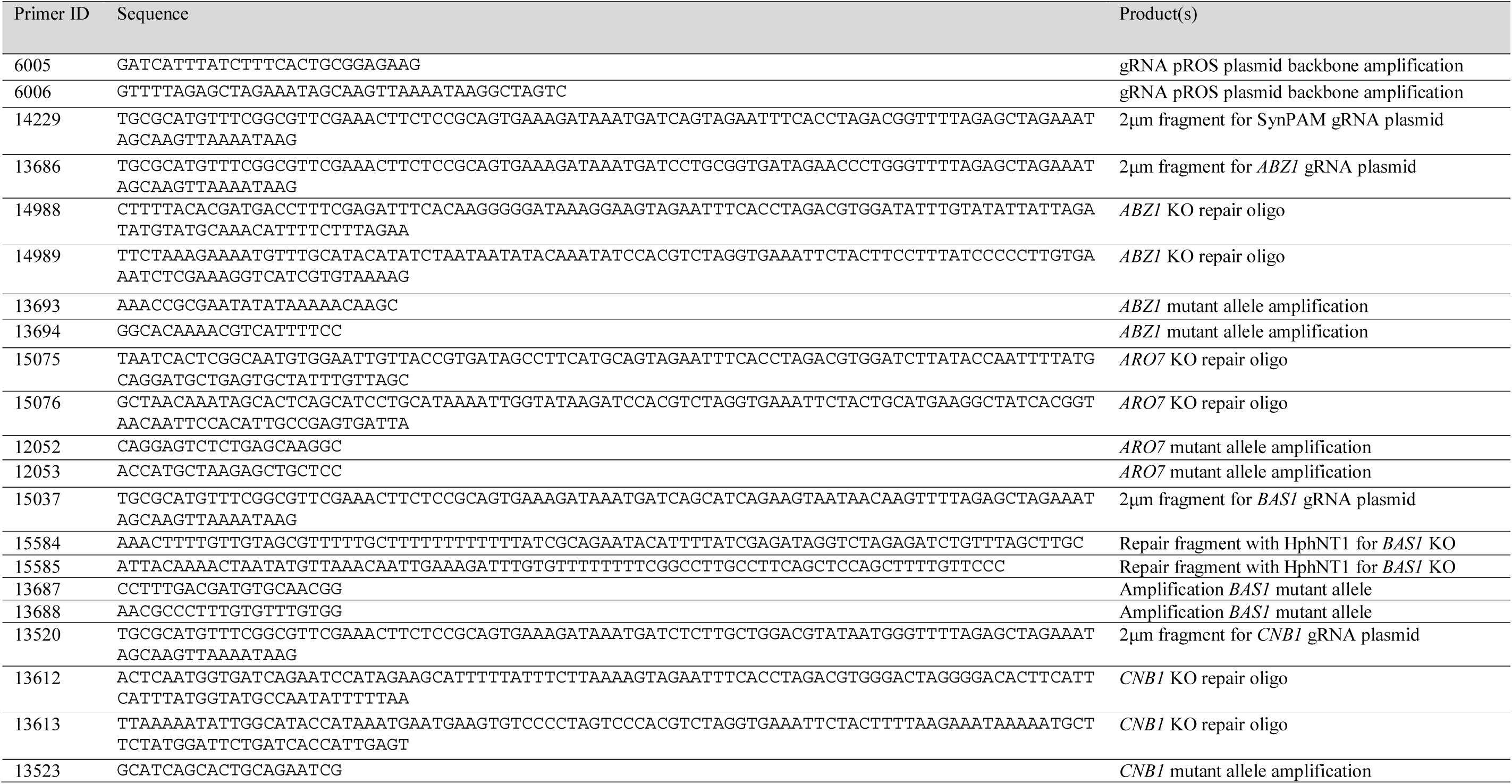
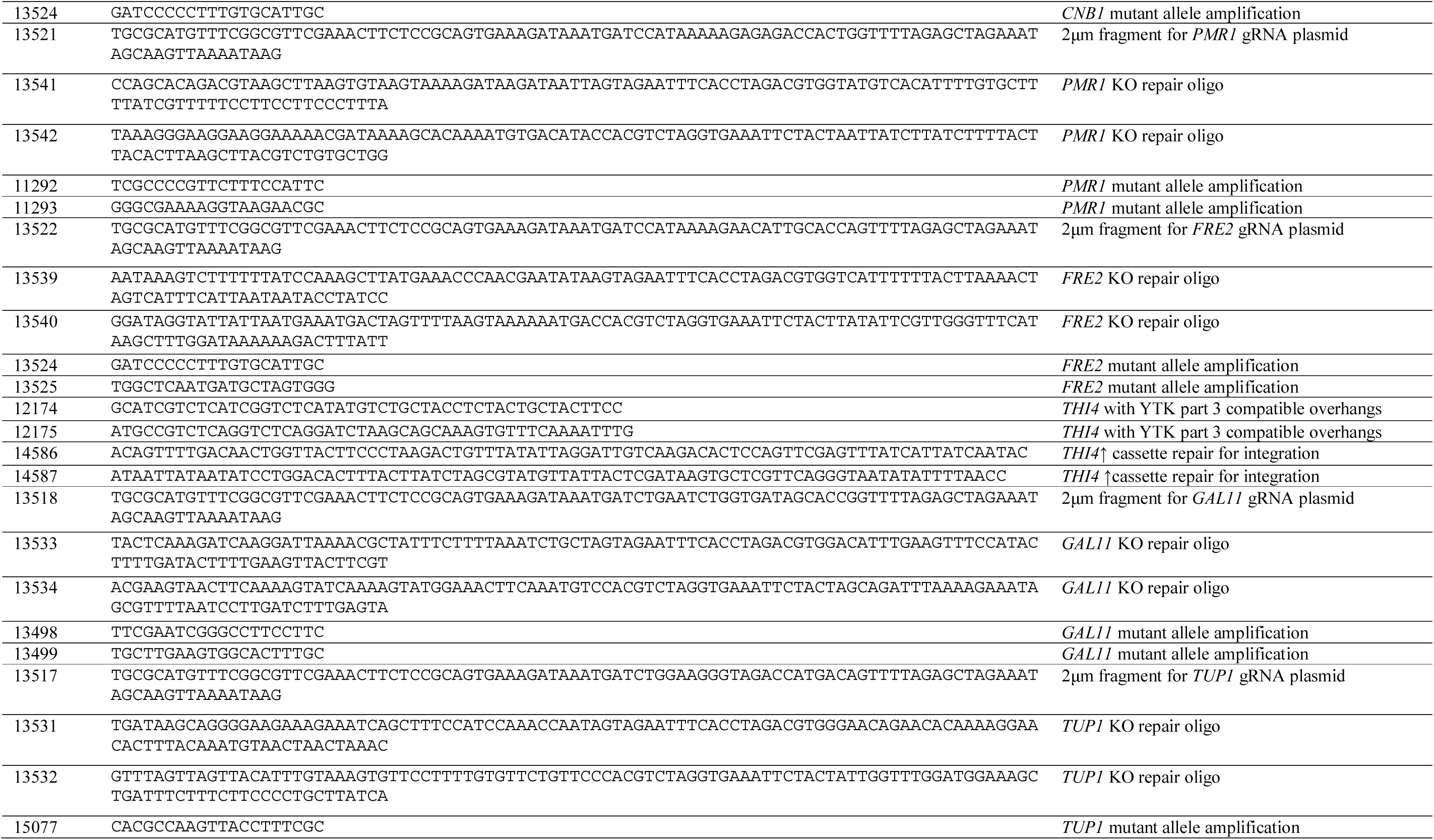
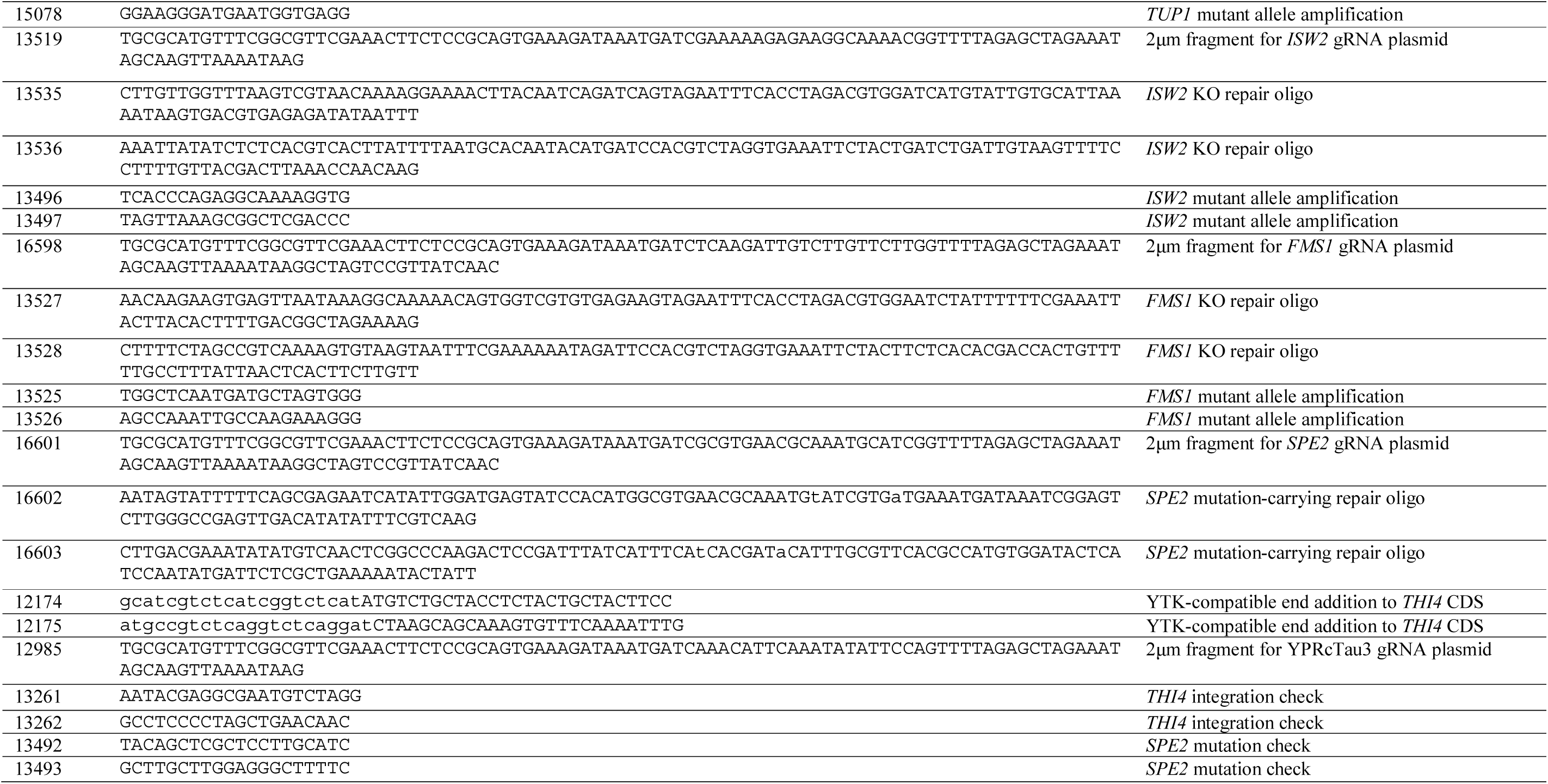
Oligonucleotide primers used in this study.

### Laboratory evolution

Laboratory evolution of *S. cerevisiae* CEN.PK113-7D for fast growth in SMD medium lacking a single vitamin was performed by sequential transfer in aerobic shake-flask batch cultures. A frozen aliquot of strain CEN.PK113-7D was inoculated in a pre-culture shake flask containing SMD medium supplemented with all vitamins. Cells were then spun down, washed twice with sterile water and used to inoculate a second shake flask containing SMD lacking one of the vitamins. The culture was then grown until stationary phase and transferred in a third shake flask containing the same fresh medium. At each transfer, 0.2 ml culture broth were transferred to 20 ml fresh medium, corresponding to about 6.7 generations in each growth cycle. The evolution experiment was performed in SMΔthiamine, SMΔpyridoxine, SMΔpantothenic acid, SMΔinositol, SMΔnicotinic acid, SMΔpABA media. Each evolution experiment was performed in triplicate. After a defined numbers of transfers, intermediate strains were stocked and characterized for the growth rate. The experiment was stopped once the target specific growth rate of 0.35 h^-1^ was reached. From each evolved population, three single colonies were then isolated and stored. The specific growth rate of these single cell lines was measured to verify that they were representative of the evolved population. The best performing isolate from each evolution line was selected for whole-genome sequencing.

### Shake flask growth experiments

For specific growth rate measurements of strains (evolved populations as well as single cell lines), an aliquot was used to inoculate a shake flask containing 100 ml of fresh medium. For specific growth rate measurements of the engineered strains, a frozen aliquot was thawed and used to inoculate a 20 ml starter culture that was then used to inoculate the 100 ml flask. An initial OD_660_ of 0.1 or 0.2 was used as a starting point. The flasks were then incubated, and growth was monitored using a 7200 Jenway Spectrometer (Jenway, Stone, United Kingdom). Specific growth rates were calculated from at least four time-points in the exponential growth phase of each culture.

### DNA sequencing

Genomic DNA of strains IMS0721, IMS0722, IMS0723, IMS0724, IMS0725, IMS0726, IMS0727, IMS0728, IMS0729, IMS0730, IMS0731, IMS0732, IMS0733, IMS0734, IMS0735, IMS0736, IMS0737, IMS0738, IMS0747, IMS0748, IMS0749, IMX2128, IMX2135, IMX2136, and IMX2137 was isolated with a Blood & Cell Culture DNA Kit with 100/G Genomics-tips (QIAGEN, Hilden, Germany) according to the manufacturers’ protocol. Illumina-based paired-end sequencing with 150-bp reads was performed on 300-bp insert libraries Novogene (Novogene (HK) Company Limited, Hong Kong) with a minimum resulting coverage of 50x. Data mapping was performed against the CEN.PK113-7D genome (22) where an extra chromosome containing the relative integration cassette was previously added. Data processing and chromosome copy number variation determinations were done as previously described (59, 69).

### Plasmids cloning

Plasmids carrying two copies of the same gRNA were cloned by *in vitro* Gibson assembly as previously described (70). In brief, an oligo carrying the 20 bp target sequence and homology to the backbone plasmid was used to amplify the fragment carrying the 2µm origin of replication sequence by using pROS13 as template. The backbone linear fragment was amplified by using primer 6005 and either pROS12 or pROS13 as template (71). The two fragments were then gel purified, combined and assembled *in vitro* using the NEBuilder HiFi DNA Assembly Master Mix (New England BioLabs, Ipswich, MA) following manufacturer’s instructions. Transformants were selected on LB plates supplemented with 100 mg/L ampicillin.

Primers 13520, 13521, 13522, 13686, 13518, 14229, 14271, 14272, 14848, 15037, 15728, 12985, 16598, 16601 were used to amplify the 2µm fragments targeting *CNB1*, *PMR1*, *FRE2*, *ABZ1*, *GAL11*, SynPAM, *TUP1*, *ISW2*, *BAS1*, hphNT1, YPRcTau3, *FMS1*, and *SPE2,* respectively. The fragment targeting *GAL11* was cloned in a pROS12 backbone yielding plasmid pUDR441. The fragment targeting *CNB1*, *PMR1*, *FRE2*, *ABZ1*, SynPAM, *TUP1*, *ISW2*, *BAS1*, hphNT1, YPRcTau3, *FMS1*, and *SPE2* were cloned in a pROS13 backbone yielding plasmids pUDR388, pUDR389, pUDR390, pUDR438, pUDR471, pUDR472, pUDR473, pUDR566, pUDR650, pUDR571, pUDR514, pUDR652, and pUDR651, respectively.

The plasmid carrying the expression cassette for *THI4* was cloned by golden gate assembly using the yeast toolkit parts (72). The *THI4* coding sequence was amplified using the primer pair 12174/12175 and CEN.PK113-7D genomic DNA as a template in order to add YTK compatible ends to the gene. The PCR product was then purified and combined together with plasmids pYTK009, pYTK056, and pYTK096 in a BsaI golden gate reaction that yielded plasmid pUDI180.

### Strain construction

Strains carrying the target mutations were all constructed starting from IMX585 expressing the Cas9 protein (71). For all strain except for IMX2290, IMX2291, IMX2289 and IMX2308, a two-steps strategy was adopted where first the target gene to be mutated was removed and replaced with a synthetic and unique 20 bp target sequence + 3 bp PAM sequence (SynPAM) and then, the synthetic target sequence was targeted and replaced with the mutant gene. In the second step where the SynPAM sequence was targeted, the mutant gene flanked by about 400 bp upstream and downstream sequences was amplified by using the evolved strain genomic DNA as template. The PCR product was then gel purified and used as repair-fragment in the transformation. This strategy yielded both intermediate strains lacking the targeted gene and final strains carrying the desired mutant gene.

In the first step, IMX585 was targeted at the gene of interest by transforming the strain with the relative pUDR plasmid. The double-strand break was then repaired by co-transforming the strain with two complementary DNA oligos carrying the SynPAM sequence flanked by 60 bp homology sequences to the targeted *locus* that were previously combined at 1:1 molar ratio, boiled for 5 minutes and annealed by cooling down the solution at room temperature on the bench.

500 ng of annealed primers pair 13612/13613, 13541/13542, 13539/13540, 14988/14989, 15075/15076, 13533/13534, 13531/13532, 13535/13536, 13527/13528 were co-transformed with 500 ng pUDR388, pUDR389, pUDR390, pUDR438, pUDR412, pUDR441, pUDR472, pUDR473, pUDR652 respectively yielding IMX1721, IMX1722, IMX1723, IMX1988, IMX1820, IMX1819, IMX1817, IMX1818, IMX2292 respectively. IMX1819 and IMX1820 transformants were selected on YPD plates with 200 mg/L hygromycin while IMX1721, IMX1722, IMX1723, IMX1988, IMX1817, IMX1818, and IMX2292 transformants were selected on YPD plates with 200 mg/L G418.

The *BAS1* knock-out strain could not be obtained with the marker-free SynPAM strategy. Therefore, the hphNT1 marker cassette was amplified by using primers 15584/15585 to add 60 bp homology flanks and pROS12 as a template. The PCR fragment was then gel purified and 500 ng were co-transformed with 500 ng pUDR592 to yield IMX2128. Transformants were selected on YPD plates with 200 mg/L G418 and 200 mg/L hygromycin.

In the second step, the SynPAM target sequence in each knock out strain was targeted for the insertion of the mutant allele. The mutant gene flanked by about 400 bp upstream and downstream sequences was amplified using the evolved strain genomic DNA as template. The PCR product was then gel purified and 500 ng were co-transformed with 500 ng of pUDR471. Primer pairs 13523/13524, 11292/11293, 13525/13526, 12052/12053, 11725/11726, 13498/13499, 13498/13499, 15077/15078, 15077/15078, 13496/13497, 13527/13528 were used to amplify the mutant alleles of *CNB1*^L82F^, *PMR1*^S104Y^, *FRE2*^T110S^, *ARO7*^L205S^, *ABZ1*^R593H^, *GAL11*^I541N^, *GAL11*^Q383Stop^*, TUP1*^V374A^, *TUP1*^Q99Stop^, *ISW2*^S181Stop^, and *FMS1*^Q33K^, respectively using IMS0747, IMS0748, IMS0748, IMS0728, IMS0727, IMS0734, IMS0735, IMS0734, IMS0735, IMS0733, IMS0736, IMS0735 genomic DNA as template, respectively. Transformants were selected on YPD plates with 200 mg/L G418, yielding IMX1985, IMX1986, IMX1987, IMX1989, IMX2057, IMX1991, IMX1992, IMX2002, IMX2003, IMX1990, and IMX2292, respectively. The *BAS1*^Q152R^, *BAS1*^D101N^, *BAS1*^S41P^ mutant alleles were amplified from IMS737, IMS738, and IMS739 genomic DNA respectively using the primer pair 13687/13688. After gel purification, 500 ng of each PCR product was co-transformed in IMX2128, together with the hphNT1 targeting plasmid pUDR650, yielding IMX2135, IMX2136, and IMX2137, respectively. The strain IMX2289 carrying the *SPE2*^A278T^ mutant allele was constructed by transforming IMX585 with the *SPE2* targeting plasmid pUDR651 together with the annealed primer pair 16602/16603 containing the desired single base change plus a synonymous mutation causing the removal of the PAM 26 sequence. After transformation, strains IMX2135, IMX2136, IMX2137, and IMX2289 were plated on YPD plates with 200 mg/L G418 for selection.

Mutant alleles found in the same evolved strains were combined in a single strain by repeating the strategy described above but this time using a mutant strain as a starting point instead of IMX585. In this way, *GAL11*, *TUP1*, and *FMS1* were deleted in IMX2002, IMX2003, and IMX2127 respectively by co-transforming the relative gRNA plasmid and the relative dsDNA oligo pair as done for the single knock out strains, yielding the intermediate strains IMX2066, IMX2110, and IMX2294 respectively. Then, the SynPAM sequence was targeted in IMX2066, IMX2110, and IMX2294 as previously described for the single mutant strains, yielding IMX2087, IMX2127, and IMX2307 respectively. IMX2043 carrying the *PMR1*^S104Y^-*FRE2*^T110S^ double mutation was constructed by co-transforming IMX1987 with pUDR390 and the linear fragment containing the *FRE2*^T110S^ mutant allele that was previously amplified as described above. The *SPE2*^A278T^ mutant allele was combined with the *GAL11*^I541N^ *TUP1*^V374A^ mutant alleles present in IMX2127 by co-transforming the strain with the *SPE2* targeting plasmid pUDR651 together with the annealed primer pair 16602/16603, yielding IMX2308. The *THI4* overexpression cassette was amplified by using pUDI180 as a template and primers 12174/12175. 500ng of gel-purified PCR product was co-transformed together with the YPRcTau3 targeting plasmid pUDR514 in IMX585 and IMX1985 yielding IMX2290 and IMX2291 respectively.

To verify the correct gene editing, single colonies were picked from each transformation plate and genomic DNA was extracted as previously described (66). The targeted *locus* was amplified by PCR and run on a 1% agarose gel. Primers pair 13523/13524, 13541/13542, 13539/13540, 15077/15078, 13496/13497, 13498/13499, 12052/12053, 13523/13524, 13541/13542, 13539/13540, 13693/13694, 12052/12053, 13496/13497, 13498/13499, 13498/13499, 15077/15078, 15077/15078, 13524/13525, 13693/13694, 13498/13499, 15077/15078, 15077/15078, 13687/1 3688, 13498/13499, 13687/13688, 13687/13688, 13687/13688, 13261/13262, 13261/13262, 13492/13493, 13525/13526, 13525/13526, 13492/13493, 13525/13526 were used to verify the correct gene editing in IMX1721, IMX1722, IMX1723, IMX1817, IMX1818, IMX1819, IMX1920, IMX1985, IMX1986, IMX1987, IMX1988, IMX1989, IMX1990, IMX1991, IMX1992, IMX2002, IMX2003, IMX2043, IMX2057, IMX2066, IMX2110, IMX2127, IMX2128, IMX2087, IMX2135, IMX2136, IMX2137 IMX2290, IMX2291, IMX2289, IMX2292, IMX2306, IMX2308, and IMX2307 respectively. To verify the presence if the single point mutations, each PCR product was purified and Sanger sequenced (Baseclear, The Netherlands). Mutations in *BAS1* could not be verified by Sanger sequencing and therefore whole-genome re-sequencing of IMX2135, IMX2136, IMX2137 was performed as explained above for the evolved single colony isolates.

After genotyping of the transformants, correct isolates were grown in 20 ml YPD in a 50 ml vented Greiner tube at 30 °C overnight by inoculating a single colony. The next day, 1 µl was transferred to a new tube containing the same amount of medium and the sample was grown overnight. The day after, each liquid culture was restreaked to single colony by plating on YPD agar plates. Plates were incubated at 30 °C overnight and the next day single colonies were patched on both YPD and YPD plus the relative antibiotic (either G428 or hygromycin) to assess which clones have lost the gRNA plasmid. One clone for each strain that had lost the plasmid was then grown in YPD and 30 %v/v glycerol was added prior to stocking samples at −80 °C.

### Data availability

The sequencing data of the evolved and of the *BAS1* deletion *Saccharomyces cerevisiae* strains were deposited at NCBI (https://www.ncbi.nlm.nih.gov/) under BioProject accession number PRJNA603441. All measurement data used to prepare the figres of the manuscript are available at the data.4TU.nl repository under the doi: 10.4121/uuid:53c9992f-d004-4d26-a3cd-789c524fe35c.

## Acknowledgments

Experiments were designed by TP, JMD and JP. Strain evolution and isolation was performed by TP. Analysis of next-generation sequencing data was performed by MvdB and TP. Reverse engineering of target mutations and phenotypical characterization of the strains was done by TP and DM. TP and JMD wrote the first version of manuscript. All authors critically read this version, provided input and approved the final version. This work has received funding from the European Union’s Horizon 2020 research and innovation program under the Marie Sklodowska-Curie grant agreement No 722287.

